# The Evolutionary History of Small RNAs in the Solanaceae

**DOI:** 10.1101/2021.05.26.445884

**Authors:** Patricia Baldrich, Sébastien Bélanger, Shuyao Kong, Suresh Pokhrel, Saleh Tamim, Chong Teng, Courtney Schiebout, Sai Guna Ranjan Gurazada, Pallavi Gupta, Parth Patel, Hamid Razifard, Mayumi Nakano, Ayush Dusia, Blake C. Meyers, Margaret H. Frank

## Abstract

The Solanaceae or “nightshade” family is an economically important group that harbors a remarkable amount of diversity. To gain a better understanding of how the unique biology of the Solanaceae relates to the family’s small RNA genomic landscape, we downloaded over 255 publicly available small RNA datasets that comprise over 2.6 billion reads of sequence data. We applied a suite of computational tools to predict and annotate two major small RNA classes: (1) microRNAs (miRNAs), typically 20-22 nt RNAs generated from a hairpin precursor and functioning in gene silencing, and (2) short interfering RNAs (siRNAs), including 24-nt heterochromatic siRNAs (hc-siRNAs) typically functioning to repress repetitive regions of the genome via RNA-directed DNA methylation, as well as secondary phased siRNAs (phasiRNAs) and trans-acting siRNAs (tasiRNAs) generated via miRNA-directed cleavage of a Pol II-derived RNA precursor. Our analyses described thousands of small RNA loci, including poorly-understood clusters of 22-nt siRNAs that accumulate during viral infection. The birth, death, expansion, and contraction of these small RNA loci are dynamic evolutionary processes that characterize the Solanaceae family. These analyses indicate that individuals within the same genus share similar small RNA landscapes, whereas comparisons between distinct genera within the Solanaceae reveal relatively few commonalities.

**ONE-SENTENCE SUMMARY:** We use over 255 publicly-available small RNA datasets to characterize the small RNA landscape for the Solanaceae family.

## INTRODUCTION

Plant genomes encode thousands of small RNA (sRNA) loci that participate in a wide variety of processes, including developmental transitions, epigenetic patterning, biotic and abiotic stress responses, and local and long-distance communication. sRNAs are 18 to 24 nucleotide (nt) non-coding sequences that are loaded into Argonaute (AGO) proteins to direct transcriptional or post-transcriptional gene silencing in a sequence-specific manner. sRNAs can be divided into different classes based on their length, biogenesis, and mode of action. This study focuses on microRNAs (miRNAs) and short interfering RNAs (siRNAs).

MicroRNAs are typically 21- or 22-nt in length and originate from a self-complementary, single-stranded (ss)RNA molecule, a Pol II product, that forms a hairpin structure. This characteristic secondary structure is typically recognized by DICER-LIKE 1 (DCL1) and processed through two sequential cleavage events into a mature double-stranded (ds)RNA duplex; each of these mature miRNAs has the potential to be loaded into Argonaute effector proteins (Axtell et al., 2011).

Small interfering RNAs (siRNAs) can be divided into multiple categories: hairpin siRNAs, which are generated in a manner similar to miRNAs, heterochromatic siRNAs (hc-siRNAs), and secondary siRNAs. hc-siRNAs are the most abundant and diverse sRNA class in plants; they are typically 24-nt in length and originate from heterochromatic regions of the genome (e.g. repetitive regions and transposable elements). hc-siRNA precursors are transcribed by RNA Pol IV, converted into dsRNA by RNA-DEPENDENT RNA POLYMERASE 2 (RDR2), and finally recognized and cleaved by DCL3, generating mature duplexes of which individual sRNAs guide sequence-specific RNA-directed DNA methylation (RdDM) (Won et al., 2014). In contrast, secondary siRNAs are generated from coding and non-coding Pol II transcripts and require a miRNA “trigger” to initiate production. Secondary siRNAs include phased siRNAs (phasiRNAs), trans-acting siRNAs (tasiRNAs), and epigenetically activated siRNAs (easiRNAs). PhasiRNA production is typically initiated by a 22-nt miRNA trigger that recognizes and directs the cleavage of ssRNA targets, which are then transformed into dsRNA by RDR6, and processed into “in phase” sRNAs via cleavage in a sequential manner by DCL4 and DCL5 (Fei et al., 2013; Chen et al., 2018b).

The evolutionary history of sRNA biogenesis pathways is complex. While the core pathways supporting the biogenesis of major sRNA classes are conserved across the land plants, lineage-specific instances of duplication, subfunctionalization, and gene loss often lead to complicated evolutionary relationships amongst AGO, DCL, and RDR family members. In the AGO family, the moss genome of *Physcomitrella patens*, for instance, contains three *AGO* genes (AGO1/4/7), while flowering plant genomes often contain 10 to 20 *AGO* family members (Axtell, 2018). The Arabidopsis genome, for example, contains ten *AGO* genes encoding proteins that can be broadly grouped into three clades: AGO1/5/10, AGO2/3/7/8, and AGO4/6/9 (Zhang et al., 2015). These individual AGO clades interact with distinct classes of sRNAs. For example, AGO1 predominantly binds 21 or 22-nt miRNAs, AGO4 binds 24-nt hc-siRNAs in the canonical RNA dependent DNA methylation (RdDM) pathway, while AGO2 binds 22-nt siRNAs that function in viral responses (Ma and Zhang, 2018). Additional AGOs have undergone further subfunctionalization in select species, for example, there are four AGO1 paralogs in rice, each of which has distinct miRNA binding affinities (Wu et al., 2009). There are also instances of *de novo* function arising within the AGO gene family. For example, maize AGO18b is a grass-specific AGO that functions the regulation of inflorescence meristem development and negatively regulates spikelet number on the tassel (Sun et al., 2018). Despite the functional importance of gene family expansion and contraction amongst key components of sRNA biogenesis machinery, relatively little characterization of the DCL, RDR, and AGO families has been carried out in the Solanaceae. Existing data focused on small subsets of Solanaceae species have characterized lineage-specific expansion events within the AGO (Liao et al., 2020), RDR, and DCL families (Esposito et al., 2018). Given the abundance of genomic resources now available for the Solanaceae, a more comprehensive, cross-species characterization of these gene family members will help resolve questions with respect to the evolution of unique aspects of sRNA biogenesis within this family.

The Solanaceae harbors a remarkable amount of morphological and metabolomic diversity. The nightshade family includes staple crops (e.g. potato and eggplant), high-value crops (e.g. tomato, pepper, petunia, and tobacco), and emerging crop systems (e.g. ground cherry), making it the third most economically important plant family in the world. Moreover, several species in the Solanaceae family serve as popular model systems for studies of biological processes pronounced within this family, including fruit ripening (Klee and Giovannoni, 2011; Shinozaki et al., 2018), plant-pathogen interactions (Goodin et al., 2015; Pombo et al., 2020), and the production of specialized metabolites (Fan et al., 2019; Leong et al., 2019). These model systems are supported by high-quality reference genomes (Mueller et al., 2005; Potato Genome Sequencing Consortium et al., 2011; Bombarely et al., 2012; Tomato Genome Consortium, 2012; Qin et al., 2014; Bombarely et al., 2016) that have facilitated the generation and analysis of genomic-scale deep sequencing datasets within the family. There are currently more than 250 Solanaceae libraries, equating to over 2.6 billion reads of publicly-available sRNA data available in the NCBI short read archive (SRA). Previous work focused on specific aspects of sRNA biology within the Solanaceae have identified unique sRNA phenomena that are associated with the diverse biology of this family. One specific case is the miR482/miR2118 superfamily in Solanaceae, which also includes miR5300 members. In eudicots, this family of 22-nt miRNAs targets the encoded sequence of the P-loop motif of NLR disease resistance proteins (Shivaprasad et al., 2012), generating a secondary siRNA cascade with implications for basal innate immunity (Zhai et al., 2011; Shivaprasad et al., 2012). Evolutionary studies indicate that this superfamily originated prior to the split of the Solanum and *Nicotiana* genera (de Vries et al., 2015), and superfamily members have even been found in Gymnosperm species, indicating that this miRNA family has deep evolutionary origins (Xia et al., 2015).

Genomic resources that support cross-species analyses in land plants have grown substantially over the last two decades. There are now more than 200 fully-sequenced genomes covering major branches of the land plant phylogeny. In addition to the availability of plant genomes, innovations in RNA library construction and sequencing technologies, as well as algorithms and bioinformatic resources, have drastically reduced experimental costs and facilitated the large-scale production of publicly-available sRNA sequence data. Two recent studies using publicly-available and newly generated sRNA datasets harvested from a broad range of land plant taxa have taken advantage of these resources and substantially improved the annotation of miRNA, phasiRNA and hc-siRNA loci in land plant genomes (<u>Lunardon et al., 2020</u>; <u>Chen et al., 2019</u>). These studies both show that siRNA loci are minimally conserved across species, while mature miRNAs tend to be conserved across lineages (<u>Chen et al., 2019</u>; <u>Lunardon et al., 2020</u>). There are numerous lineages- or species-specific miRNA sequences which correspond to young miRNAs (<u>Cuperus et al., 2011</u>; <u>Chávez Montes et al., 2014</u>; <u>Lunardon et al., 2020</u>). Although large-scale studies produce a massive amount of data and describe major evolutionary trends, they are often unable to capture lineage- and species-specific phenomena.

In this study, we applied a suite of computational tools to a large collection of publicly-available sRNA libraries to investigate the unique aspects of miRNAs and siRNA biology within the Solanaceae. We describe thousands of sRNA loci, some of which are unique to the Solanaceae. For example, we found that an annotated, Solanaceae-specific miRNA family is actually a MITE-derived siRNA, and we observed family-wide expression of *DCL2*-derived 21-nt phasiRNAs. Our analyses also revealed aspects of sRNA biology that are only conserved at the sub-family level, including a lineage-specific expansion of previously unidentified miRNAs in *Nicotiana*, a distinct split in phasiRNA production between the *Solanum*/*Capsicum* and *Nicotiana*/*Petunia* branches of the family, and distinct clusters of 22-nt sRNAs that are expressed in response to viral infection in tomato. Overall, our analyses indicate that individuals within the same genus share similar sRNA landscapes, whereas comparisons between distinct genera within the Solanaceae reveal relatively few commonalities.

## RESULTS

### Evolution of small RNA biogenesis genes in the Solanaceae

sRNA biogenesis and function are regulated by proteins in the Nuclear RNA Polymerase (NRP), RNA-dependent RNA polymerase (RDR), Double-stranded RNA binding protein (DRB), Dicer-Like protein (DCL) and Argonaute protein (AGO) families. To describe and interpret sRNA biogenesis and function in the Solanaceae, we performed a phylogenetic analysis of these proteins encoded in genomes of *Arabidopsis thaliana*, two Convolvulaceae outgroups (*Ipomoea trifida* and *Ipomoea triloba*) and seven Solanaceae species (*Nicotiana benthamiana*, *N. tabacum*, *Petunia axillaris*, *P. inflata*, *Capsicum annuum*, *Solanum lycopersicum* and *S. tuberosum*). In Supplemental Table 1 we summarize the number of protein genes encoded for *NRP*, *RDR*, *DCL*, *DRB* and *AGO* gene families in each species. Two observations emerged from this table. First, some gene families expanded when comparing Arabidopsis to members of the Convolvulaceae and/or Solanaceae families. For instance, the number of genes expanded in the AGO family comparing *A. thaliana* (10 *AGOs*) to Convolvulaceae (13 *AGOs*) or Solanaceae for the genus *Petunia* (12 or 13 *AGOs*) and the genus *Solanum* (13 or 15 *AGOs*). A second observation was the appearance of missing gene annotations for specific gene families in certain species (e.g. *NRP* genes encoded in tomato and potato). Thus, we chose to focus on *AGO* and *DCL* gene families in all species. All annotated genes are detailed in Supplemental Table 1. Their evolutionary relationships are visualized in Figure 1 and Supplemental Figure 1.

**Figure 1:**
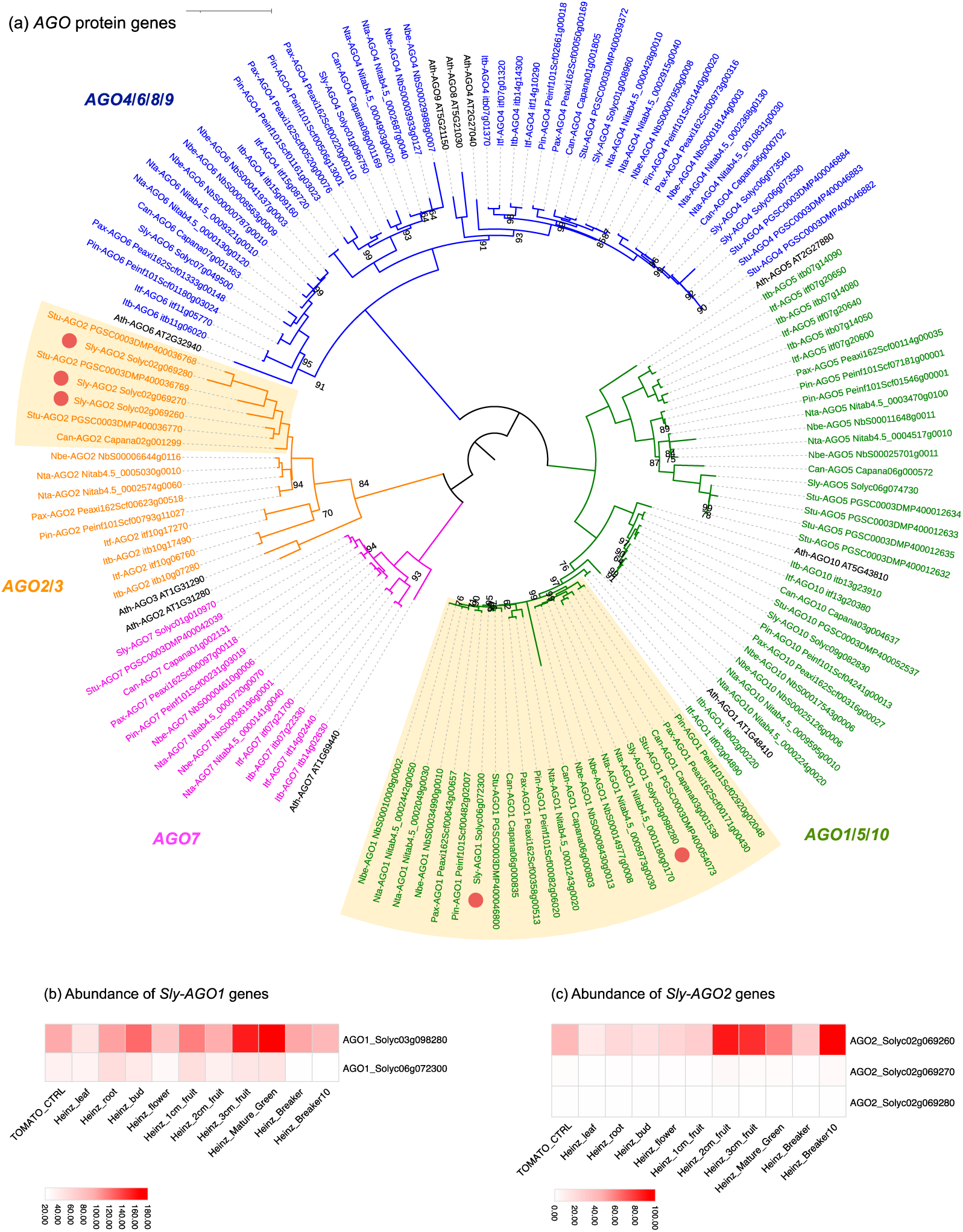
Solanaceae-specific diversification of the *AGO* family highlights expansion and subfunctionalization of subfamily members. Phylogenetic tree of proteins encoded by the *AGO* gene family in the Solanaceae (A). Clades of *AGO* subfamily members are drawn with distinct colors. Yellow boxes highlight groups of *AGO* genes that have expanded in Solanaceae. Red circles indicate *AGO* homologs in tomatoes. Heatmaps show the relative expression of *AGO1* and *AGO2* from public data (B). Orthologous protein groups were identified using OrthoFinder and SonicParanoid. Protein alignments and phylogeny analysis were completed using MUSCLE and IQ-TREE. Phylogenetic tree was illustrated using iTOL.

AGO proteins are required to load sRNAs and to catalyze their functional activity. AGO proteins can either trigger the biogenesis of sRNAs, through the miRNA-directed cleavage of *TAS*/*PHAS* transcripts, or catalyze sRNA regulatory activity through transcriptional or post-transcriptional regulation. Our phylogenetic analysis identified an expansion of specific and distinct *AGO* genes for Convolvulaceae and Solanaceae species when compared to *Arabidopsis*. For example, we identified a Convolvulaceae-specific expansion for *AGO5* (to three copies) and *AGO7* (to two copies) that did not occur in the Solanaceae (Figure 1; Supplemental Table 1). We also identified differential *AGO* gene family expansion and loss across the Solanaceae. For instance, we expected to find two copies of *AGO2/AGO3* (based on gene family membership in *A. thaliana*) but we only identified one homolog in *Capsicum*, *Nicotiana ssp.* And *Petunia ssp.,* and three in *Solanum ssp.* (Figure 1; Supplemental Table 1). A common feature that we observed across the Solanaceae was a family-wide expansion of *AGO1*; depending on the particular species, we identified two to three *AGO1* paralogs within the Solanaceae (Figure 1; Supplemental Table 1). To test whether this *AGO1* expansion coincides with evidence for subfunctionalization, we downloaded and visualized public expression data for the two *AGO1* paralogs we identified in *Solanum lycopersicum* (Figure 1B; http://bar.utoronto.ca/eplant_tomato/). Interestingly, we observed a distinct expression pattern of both copies with one mainly accumulating during vegetative development (Solyc06g072300), while the other mainly accumulated in fruits during reproductive development (Solyc03g098280). This observation suggests that indeed, these *AGO1* paralogs are in the process of subfunctionalization in tomato. We also discovered that *AGO2* expanded into triplicate copies that are located in tandem on chromosome 2 in tomato (Figure 1A). Existing public data indicates that only one of the three *AGO2* copies is expressed, and this is predominantly during fruit development (Figure 1C). In brief, we discovered lineage-specific expansions of *AGO* subfamily members in the Convolvulaceae and Solanaceae. Public expression data for tomato indicates that these *AGO1* and *AGO2* paralogs may have subfunctionalized following duplication.

We annotated additional sRNA biogenesis genes, including *NRP*, *RDR*, *DRB* and *DCL* (Figure 2; Supplemental Figure 1; Supplemental Table 1), and discovered a subfamily expansion of *DCL2* in the *Solanum* (tomato and potato) and *Capsicum* (pepper) branch of the Solanaceae. For all three species, *DCL2* expanded through a tandem duplication on chromosome 11 (annotated as chromosome 12 for pepper). In tomato, *DCL2* underwent additional duplication events, expanding to three tandem duplicates on chromosome 11 and additional copy on chromosome 6 (Figure 2). To gain insights into the impact of gene expansion on function, we downloaded public expression data for tomato, and found that the *DCL2* copy on chromosome 6 (Solyc06g048960) is broadly throughout the whole plant and the three paralogs on chromosome 11 (Solyc11g008520; Solyc11g008530; Solyc11g008540) are specifically enriched in flowers and fruits (Figure 2B). We also observed much higher expression levels for two of the *DCL2* paralogs on chromosome 11 (Solyc11g008530; Solyc11g008540) relative to the broadly expressed copy of *DCL2* on chromosome 6 (Solyc06g048960). Together, these observations highlight a Solanaceae-specific expansion and putative subfunctionalization of *DCL2*. See additional details about *DCL2* in our phasiRNA analysis, below.

**Figure 2:**
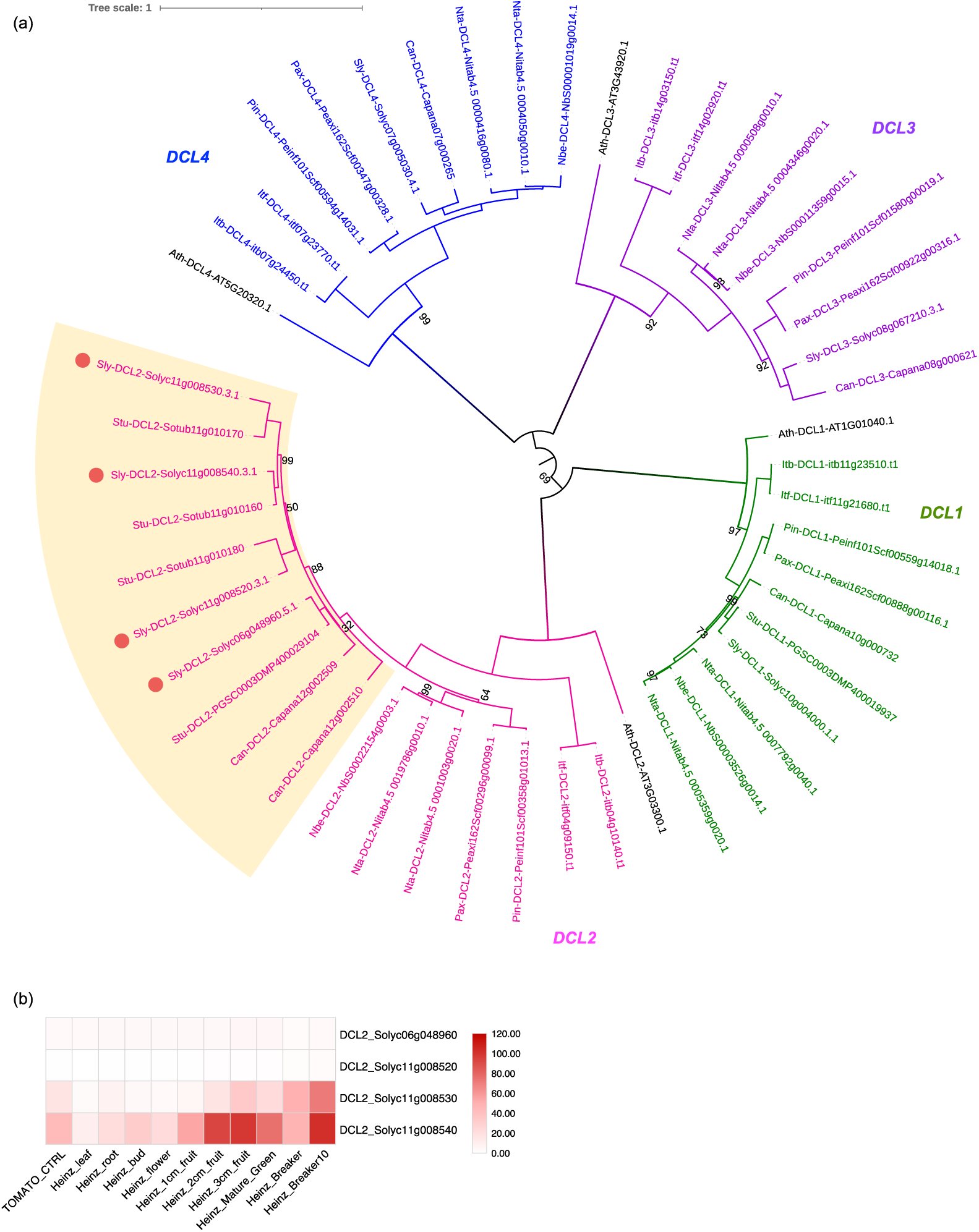
Phylogenetic analysis of DCL family members encoded in the Solanaceae highlights a *Solanum*/*Capsicum*-specific expansion of DCL2. Phylogenetic tree of protein sequences indicating *DCL* subfamily members in the Solanaceae (A). Subclades of the *DCL* family are drawn with distinct colors and yellow boxes highlight groups of *DCL2* that expanded in *Capsicum* and *Solanum* genera. Red circles indicate tomato paralogs for *DCL2*. Heatmaps show the relative abundance of these tomato *DCL2* paralogs in diverse samples (B). Protein orthologous groups were identified using OrthoFinder and SonicParanoid. Protein alignments and phylogeny analysis were done using MUSCLE and IQ-TREE. Phylogenetic tree was visualized using iTOL.

### Mining publicly-available Solanaceae sRNA data

We selected seven core species in the Solanaceae (tomato, potato, pepper, ‘petunia_ax’ *Petunia axilaris*, ‘petunia_in’ *P. inflata*, tobacco, ‘benthi’ [*Nicotiana benthamiana*]) for our cross-species analyses of sRNA loci. These species were selected based on the availability of fully sequenced genome assemblies and publicly-available sRNA data. Notably, we excluded eggplant from this study due to the paucity of publicly-available *S. melongena* sequence data. In total, we analyzed over 2.6 billion sRNA reads from 255 different libraries. By far, tomato and potato were the most extensively sampled, having 84 and 73 high-quality publicly available libraries, respectively, while tobacco was the least sampled species (represented by only 9 libraries on the NCBI SRA; Supplemental Dataset 1). The depth of sequencing also varied quite substantially across libraries. This was largely due to the differences in sequencing technologies. In line with previous studies, we counterbalanced this heterogeneity in sequence depth by merging all high-quality libraries together and treating them as independent reference sets (Lunardon et al., 2020). Most of these reference sets included organ atlas samples that represent expression patterns from diverse organ systems and developmental stages. The total number of genome aligned reads for each species varied from just over 70-million reads for *Petunia axillaris* and tobacco to more than 650 million reads for tomato (Supplemental Dataset 1).

### MicroRNA predictions indicate an expansion of novel miRNAs in *Nicotiana* species

We identified previously annotated and novel miRNAs across the Solanaceae using two software programs: ShortStack and miR-PREFeR (Axtell, 2013; Lei and Sun, 2014). While ShortStack was more conservative, and predicted fewer miRNAs in each species, our miR-PREFeR datasets included some less reliable miRNA predictions. To maximize our miRNA predictions while maintaining high-quality standards, we applied a post-prediction filter that used previously published criteria for defining miRNAs, including a minimum free energy of < -0.2 kcal/mol/nucleotide for precursor folding and the expression of both miRNA/miRNA* strands of the miRNA duplex (Axtell and Meyers, 2018). After filtering, we combined both software predictions into a consensus set of miRNAs with assigned annotations based on ≥ 85% sequence similarity to miRBase v22 (Supplemental Dataset 2). Overall, our analyses predicted between 124 and 440 miRNAs across the Solanaceous species (Figure 3; Supplemental Dataset 2). The depth of sequence data did not correlate strongly with the number of miRNAs that we predicted for a given species. For example, pepper (330) and benthi (440) had the highest number of predicted miRNAs by far, even though these species are only supported by a moderate depth of sequence data relative to tomato, which only had 194 predicted miRNAs (Supplemental Dataset 2). This indicates that our miRNA predictions were generally not constrained by sampling.

**Figure 3:**
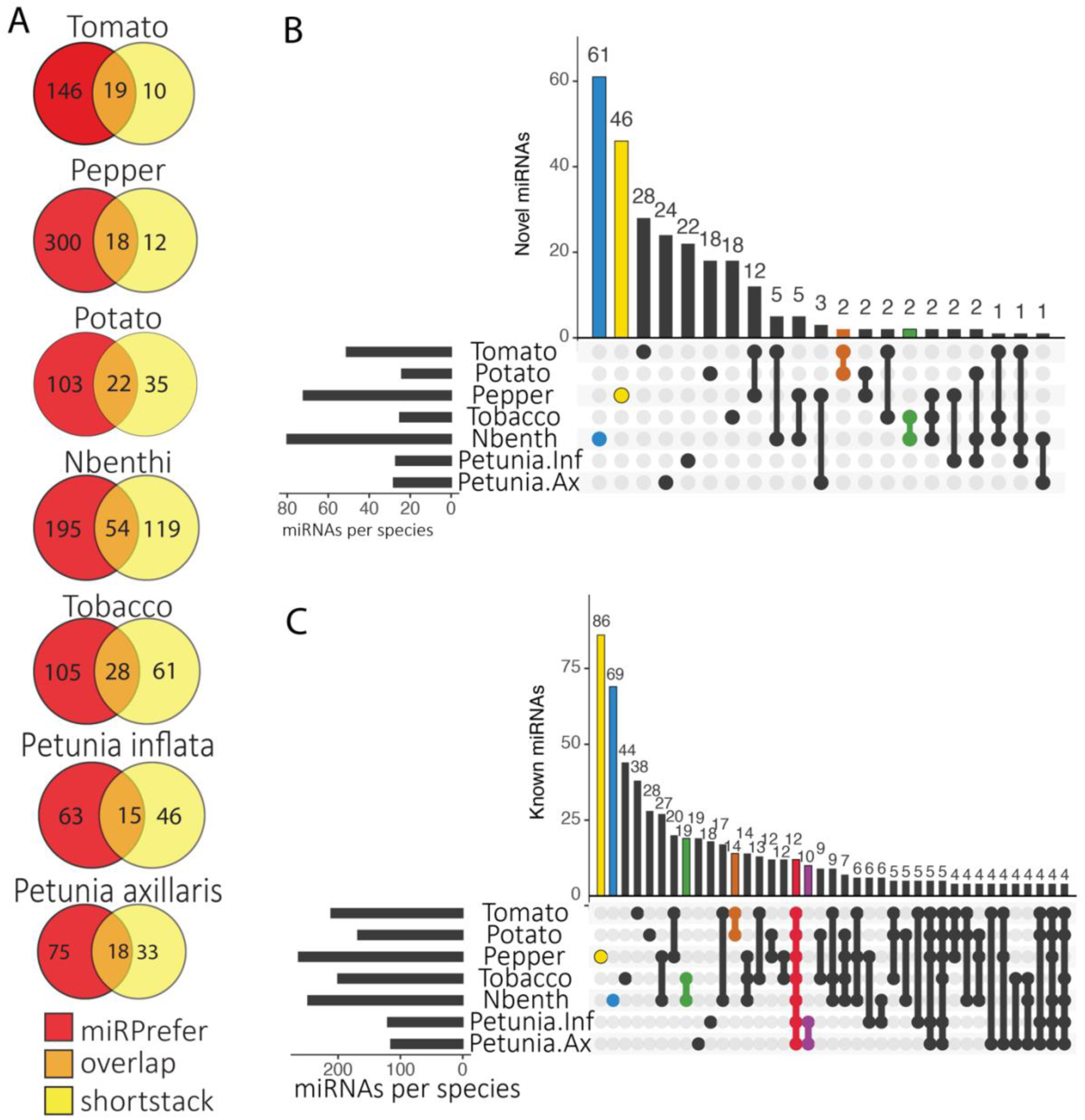
MicroRNA predictions across Solanaceae support a lineage-specific expansion of miRNAs. Venn diagrams of total miRNAs predicted by miRPrefer (red), Shortstack (yellow), and both programs (orange) for each species (A). Upset plots showing novel (B) and known (C) miRNAs that are shared across Solanaceae species. For each Upset plot: the bottom left panel shows the entire size of each set as a horizontal histogram, the bottom right panel shows the intersection matrix, and the upper right panel shows the size of each combination as a vertical histogram. The red line marks miRNAs common to all species, the purple line marks miRNAs common to both *Petunia* species, the orange line marks miRNAs common to both *Solanum* species, the green line marks miRNAs common to both *Nicotiana* species, and the blue and yellow lines mark benthi- and pepper-specific miRNAs, respectively.

We were surprised to find more than 400 predicted miRNAs in the benthi genome. To gain higher confidence in these benthi predictions, we manually filtered the benthi miRNA precursors based on the folding and expression of their predicted precursors using SructVis (Supplemental Datasets 2 & 3; https://github.com/MikeAxtell/strucVis). Even after manual filtering, we had a substantial number of predicted miRNAs for benthi (368), indicating that these predictions reflect a real lineage-specific miRNA expansion in *Nicotiana* (Figure 3). In agreement with this conclusion, we also identified a relatively high number of miRNAs in tobacco (194), despite having the lowest depth of sequence data for this species. Notably, less than half of the miRNAs that we identified in either of the two *Nicotiana* genomes could be annotated using miRBase, indicating that miRNA expansion in *Nicotiana* was likely driven by the evolution of “young” lineage-specific miRNAs. To learn how evolutionary age of these *Nicotiana*-specific miRNAs influences function, we performed miRNA target predictions with the benthi miRNA dataset (Supplemental dataset 4). We found that the more recently evolved, benthi specific miRNAs were predicted to target fewer mRNAs (33.5 mRNAs) than the miRBase-conserved miRNAs in benthi (55.6 mRNAs) (Supplemental Dataset 4), and that average penalty scores for benthi-specific miRNAs were slightly higher (3.6) than the miRBase-conserved miRNAs (3.5). Higher scores equate to a less reliable miRNA target match (Kakrana et al., 2015), therefore the benthi-specific miRNAs tend to have less reliable target predictions. Overall, this analysis indicates that more recently evolved *Nicotiana* miRNAs have fewer and less reliable targets, which may equate to more limited functions.

We also characterized a substantial expansion of predicted miRNAs in pepper (330), which is congruent with previous miRNA predictions for this species (Seo et al., 2018; Taller et al., 2018). In contrast, we predicted relatively few miRNAs for both of the petunia genomes (124 - *P. inflata* and 126 - *P. axillaris*). This reduced number of predicted miRNAs in petunia may simply reflect a lack of sample diversity for these species; publicly-available sRNA-seq datasets for petunia are predominantly composed of floral samples (Supplemental Dataset 1). Indeed, a previous analysis using combined miRNA prediction software and a homology based search for known miRNAs characterized about 140 miRNAs for both petunia genomes, indicating that our *de novo* predictions on their own are missing putative miRNA loci. Increasing the diversity of sRNA samples would likely improve miRNA annotations for this genus (Bombarely et al., 2016). Finally, we only identified a moderate number of predicted miRNAs in the Solanum genus (potato = 160, tomato = 194), despite having a substantial amount of sequence data (>80 sRNA libraries) for tomato. Solanum is by far the best characterized genus in the Solanaceae with regard to sRNA biology. Given that our predictions for this genus are similar to those from previous studies, we are likely approaching a consistent set of Solanum-specific miRNA predictions (Tomato Genome Consortium, 2012; Zhang et al., 2013; Cardoso et al., 2018; Zuo et al., 2020; Deng et al., 2021).

### miRNA families are conserved within genera

To gain insight into miRNA conservation across Solanaceous species, we collapsed all predicted miRNAs into clusters using a 75% sequence similarity cutoff, and annotated the clusters with miRBase (v22 all land plant miRNAs (Kozomara et al., 2019);Supplemental Dataset 5). This resulted in 911 clusters; 392 of which are present in two or more species, 172 that are conserved at the genus level (i.e. *Solanum*-conserved, *Petunia*-conserved, and *Nicotiana*-conserved), and only 12 that are conserved across all seven species in this study (Supplemental Dataset 5; Figure 3B-C). The 12 conserved clusters can be further condensed into a handful of known miRNA families based on annotations that are broadly conserved across the land plants (e.g. - miR156/157, miR164, miR169/miR399, miR171, miR399, miR165/166/168, and miR172, miR482/miR2118 (Axtell and Meyers, 2018). We designated clusters that could not be annotated with miRBase as “novel.” The majority of these novel clusters appear to be newly-evolved, species-specific miRNAs, while the majority of clusters that are conserved across multiple species (present across ≥ 3 species) aligned with annotated miRNAs in miRBase. In agreement with previous studies showing that lineage-specific miRNAs tend to be weakly expressed relative to conserved miRNAs, we found that the expression of conserved miRNAs was significantly higher on average relative to novel miRNAs (Supplemental Figure 2; benthi p-value = 0.00057903, pepper p-value = 1.98548E-08, potato p-value = 1.97921E-06, tomato p-value = 2.27413E-07, petunia p-value = 2.39254E-05, tobacco p-value = 5.89638E-08; (Axtell et al., 2007)).

### Solanaceae-specific miRNA family miR5303 is derived from MITE loci

Among the most conserved miRNA groupings, we identified three clusters containing a Solanaceae-specific miRNA, miR5303 (Gu et al., 2014). With the exception of petunia, the miR5303 family was present in relatively high copy numbers across the Solanaceous genomes (7 to 11 miRNAs in each species; Supplemental Dataset 5; Supplemental Figure 3). It is rare for miRNAs to be present in such high copy numbers; however, our findings are congruent with a previous study that identified a high number of miR5303 copies in tomato, potato, eggplant, and pepper (Gu et al., 2014). To correctly identify miR5303 family members, we first aligned all miR5303 variants in miRBase v22 (Figure 4), and found that most of the variants are 24-nt in length, which again, is atypical for a bonafide miRNA family (Meyers et al 2008; Fig 4A). We identified nine miR5303 variants that are 21-nt in length in our tomato predictions that have ≥ 75% similarity with miR5303 miRBase entries, and found that six of these tomato miRNAs clustered into a family containing a conserved 5’ - UUUUG consensus sequence (highlighted by yellow rectangles in Fig 4B). To further characterize the function of this family, we performed a miRNA target prediction analysis using all coding sequences in the tomato genome as putative targets. We found that miR5303 family members are predicted to target thousands of genes in the tomato genome, which again, is unusual for a miRNA family (Supplemental Dataset 6). Given the presence of 24 nucleotide mature miRNA variants that are reminiscent of 24-nt siRNAs, and the perfect/near perfect folding of miR5303 precursor sequences that are often formed by inverted repeats (Figure 4C), we decided to test whether miR5303 actually originates from miniature inverted-repeat transposable elements (MITEs) in Solanaceae genomes. Notably, MITEs exist in high copy numbers, can form hairpin sequences that are structurally similar to miRNAs, and are known to generate 24-nt siRNAs (Kuang et al., 2009). To test whether miR5303 family members originated from MITEs, we blasted precursors of miR5303 from tomato against a plant MITE database (Chen et al, 2013) and found that they align perfectly with annotated MITEs, with the miR5303 mature miRNA variants mapping to the terminal inverted repeat (TIR) ends of the MITEs (Figure 4D). We therefore conclude that miR5303 is not a miRNA, but a MITE.

**Figure 4:**
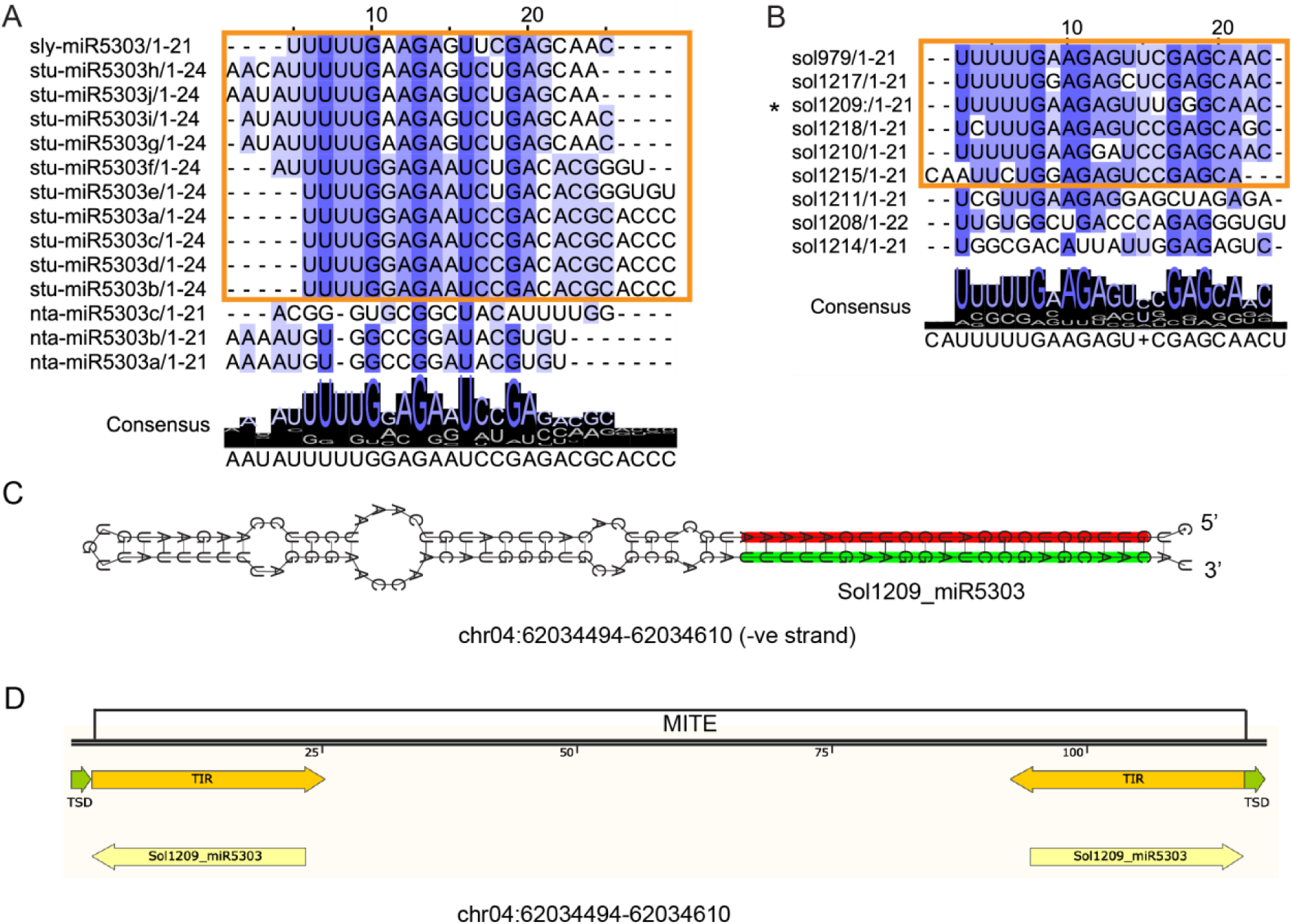
miR5303 aligns to MITE terminal inverted repeat (TIR) sequences. The previously-annotated Solanaceae-specific miRNA family, miR5303, exhibits many features that disqualify it as a bonafide miRNA. The majority of the miR5303 miRBase entries include mature sequences that are 24-nt in length, indicating that these sRNAs are likely derived from siRNA producing loci (A). miR5303 miRBase entries are highly variable in their sequence; we identified a consensus sequence to isolate true miR5303 family members (highlighted in the yellow box and shown in Supplemental Figure 3) (A). We filtered out miRNAs > 22-nt in length; however, our predicted miR5303 family members that are 21 or 22 nt follow the same 5’ consensus sequence (UUUUG) as the miRBase entries (B). The predicted precursors of miR5303 family members have near perfect hairpin folding, indicating that these loci are likely inverted repeats, rather than real miRNA precursors (C), and BLAST alignment of miR5303 family members to the plant MITE database produces a perfect alignment between the miR5303 species and MITE TIR sequences (D).

### Identification of 21-nt *PHAS* loci support genus-level conservation of phasiRNA biology

We applied two bioinformatic tools: PHASIS (Kakrana *et al*. 2017) and ShortStack (Axtell 2013), to assemble a list of 21-nt *PHAS* loci across the Solanaceae. To classify coding versus non-coding phasiRNAs we merged PHAS locus coordinates with gene annotation files for each species. Some of the lesser studied Solanaceae genomes have incomplete annotations which resulted in an overprediction of non-coding loci for petunia, pepper, and benthi. To correct this imbalance, we used BLAST to align all non-coding *PHAS* loci to the NCBI nr database (Altschul et al., 1990), and manually annotated loci that aligned to protein-coding, rRNA, and non-coding RNA sequences (Figure 5; Supplemental Dataset 7). *PHAS* loci are typically triggered by 22-nt miRNAs that have a 5’ Uracil bias (Chen et al., 2010; Cuperus et al., 2010). To identify putative triggers for 21-nt phasiRNA production, we ran target predictions using our list of mature 22-nt miRNAs plus miR390 as triggers and predicted *PHAS* loci as targets (Supplemental Dataset 8).

**Figure 5:**
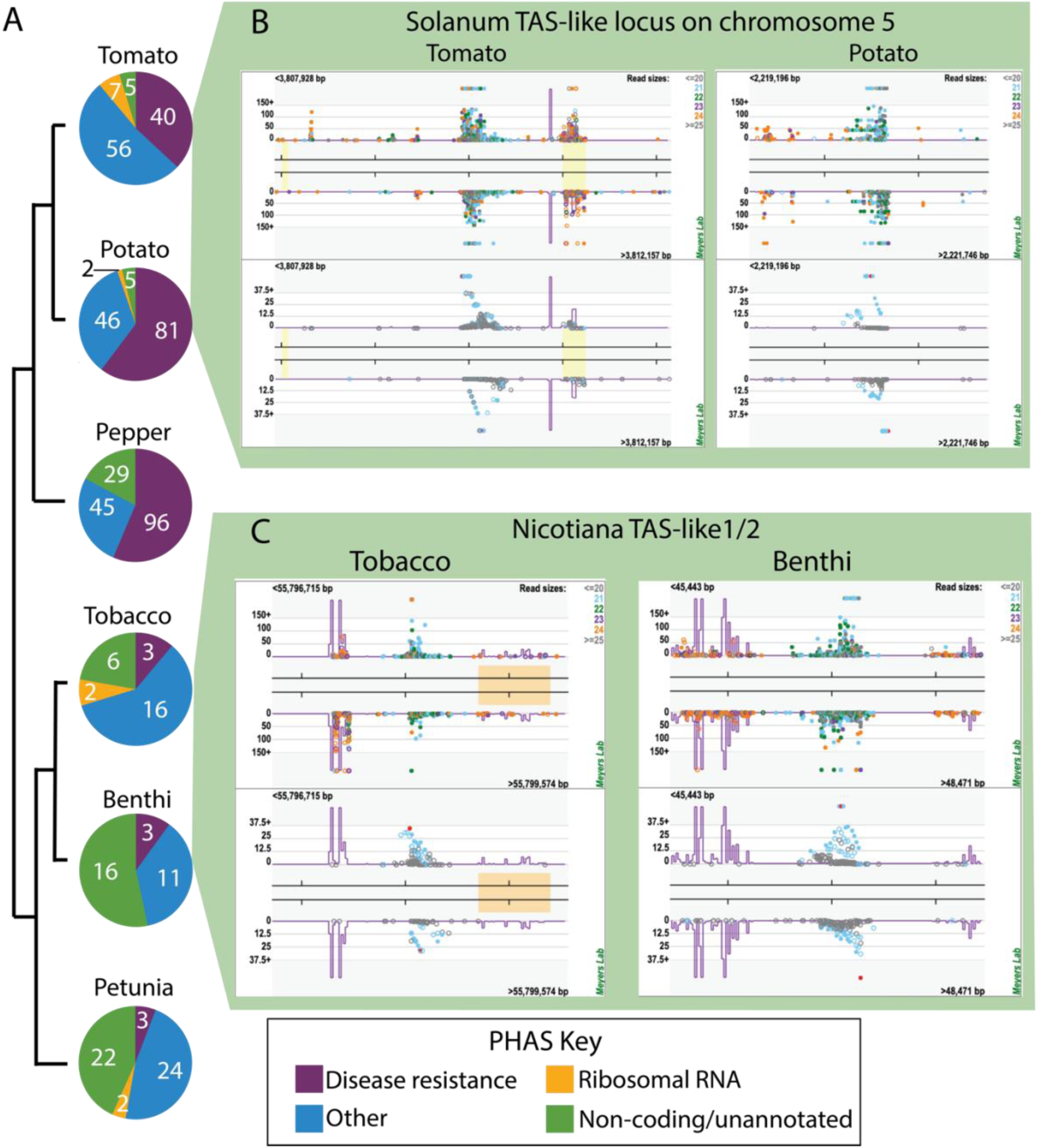
Genus-specific trends of 21-nt *PHAS* loci within the Solanaceae. *Solanum* (tomato and potato) and *Capsicum* (pepper) genera produce 2-3 times more PHAS loci and predominantly express disease resistance-related coding loci, relative to *Nicotiana* (tobacco and benthi) and *Petunia* genera (A). The raw number of loci for each category in (A) is shown in white. All of the species express non-coding PHAS loci (green). A previously unidentified non-coding TAS-like locus is shown for on chromosome 5 in tomato (left) and potato (right) (B), and a previously identified non-coding *Nicotiana-*conserved TAS-like 1/2 locus is plotted for tobacco (left) and benthi (right) (C).

PhasiRNAs are predominantly generated from transcripts of genes encoding disease resistance-related NB-LRRs, pentatricopeptide repeat (PPR) proteins, MYB transcription factors, and trans-acting siRNA (*TAS*) loci (Liu et al., 2020; Shivaprasad *et al*. 2012). We identified 2-3 times more *PHAS* loci in *Solanum* (tomato and potato) and *Capsicum* (pepper) species than *Nicotiana* and *Petunia* (Figure 5). The majority of these *Solanum*/*Capsicum* loci were derived from disease resistance-related genes that are putatively triggered by the miR482/miR2118 superfamily (Figure 5A; Supplemental Dataset 8). We also characterized an *NB-LRR* pseudogene, *TAS5*, which was recently shown to function in pathogen response in tomato (Canto-Pastor et al., 2019). In contrast, the *Nicotiana* and *Petunia* genomes expressed relatively few disease resistance related phasiRNAs, and likewise, had very few loci with predicted miR482/miR2118 target sites (Figure 5A; Supplemental Dataset 8). We also found a substantial number of non-coding loci that we decided to examine in further detail with the goal of uncovering novel *TAS* loci within the Solanaceae. As a quality control, we confirmed that *TAS3*, a deeply-conserved non-coding phasiRNA, was detected in all of our species datasets (Supplemental Dataset 7; (Xia et al., 2013)). Most notably, we discovered a previously uncharacterized, non-coding locus in both tomato and potato (Figure 5B). This locus is >80% conserved between the two species and generates highly abundant 21-nt siRNAs, some of which putatively target genes encoding MYB transcription factors (Supplemental Figure 4). We performed a BLAST alignment between the tomato *PHAS* locus and all other non-coding *PHAS* loci to determine whether this locus exists outside of Solanum. This search failed to uncover any significant alignments, indicating that this novel phasiRNA evolved after Solanum split from other genera in the Solanaceae (Särkinen et al., 2013).

As mentioned above, PPR-derived *PHAS* loci are abundantly expressed in other species. In our analysis, we only identified two PPR-related *PHAS* loci in the Solanaceae: one in tomato (Solyc10g076450) and one in potato (PGSC0003DMG400020556). We did, however, find strong evidence for another, well-described siRNA mechanism for the transcriptional regulation of PPR genes, in which non-coding *TAS-like-1/2* (*TASL1/2*) loci are triggered by miR7122, generating 21-nt products that in turn target, and likely silence, PPR genes (Xia et al., 2013; Figure 5C). We identified *TASL1/2* and its associated miR7122 trigger in both *Nicotiana* species (benthi and tobacco). As demonstrated in other species, major 21-nt products from these *TASL* loci target PPR genes (Supplemental Figure 4A-F). Surprisingly, we found no evidence for *TASL1/2* and its miR7122 trigger in other Solanaceae genera. Given the conservation of this locus in other eudicots outside of the Solanaceae family, we hypothesize that miR7122-*TASL1/2* is conserved in *Nicotiana* and was independently lost from the other genera in the nightshade family (Xia et al., 2013).

To investigate whether *PHAS* loci are generated from orthologous genes across species, we constructed gene orthogroups and searched for conserved *PHAS* loci within these orthogroupings (Supplemental Dataset 9). Overall, this analysis yielded surprisingly few orthologous *PHAS* loci within the Solanaceae. Following up on our earlier analysis of *DCL2* duplication within sub-groups of the Solanaceae, we also discovered that *DICER-LIKE2* (*DCL2*) is broadly expressed as a phasiRNA-producing locus that is present in all of the species in our study (Figure 6). Interestingly, this *PHAS* locus has been previously reported in the Fabaceae, where it is triggered by different miRNAs in medicago and soybean, indicating convergent evolution of phasiRNA production at *DCL2* (Zhai et al., 2011). A *DCL2 PHAS* locus has also been identified in tomato (Canto-Pastor et al. 2019); however, this is the first instance of its widespread presence across the nightshade family. As reported earlier in the results, *DCL2* has expanded through tandem duplications that are shared in the tomato, potato, and pepper genomes (Figure 2). We found the genomic regions containing these tandem *DCL2* paralogs to be structurally conserved on *Solanum* chr11/*Capsicum* chr12 (Figure 6A). Recently, a DCL2-dependent miRNA (miR6026) was validated as a trigger for phasiRNAs derived from the *DCL2* locus in tomato (Wang et al., 2018), creating a negative feedback loop on *DCL2*. To investigate whether the same miRNA trigger initiates *DCL2* phasiRNA production in the rest of the Solanaceae, or, as in the Fabaceae (Zhai et al., 2011), independent triggers evolved at this locus, we looked for shared miRNA-*PHAS* locus interactions. We identified a conserved target site with a strong target score for miR10533 in tomato, potato, pepper, and petunia (Figure 6B). Like miR6026, miR10533 is also dependent on DCL2 for its biogenesis (Wang et al., 2018). However, unlike in the Fabaceae, where phasiRNA production proceeds immediately after the miRNA target site (Zhai et al., 2011), *DCL2* phasiRNAs are produced both upstream and downstream of the miR10533/miR6026 target sites in the Solanaceae (Figure 6A). This imprecise initiation of phasing either indicates that there are additional miRNA triggers for this locus, or we are seeing the product of trans-triggering from 22-nt siRNAs that are generated by neighboring *DCL2* paralogs (Figure 6).

**Figure 6:**
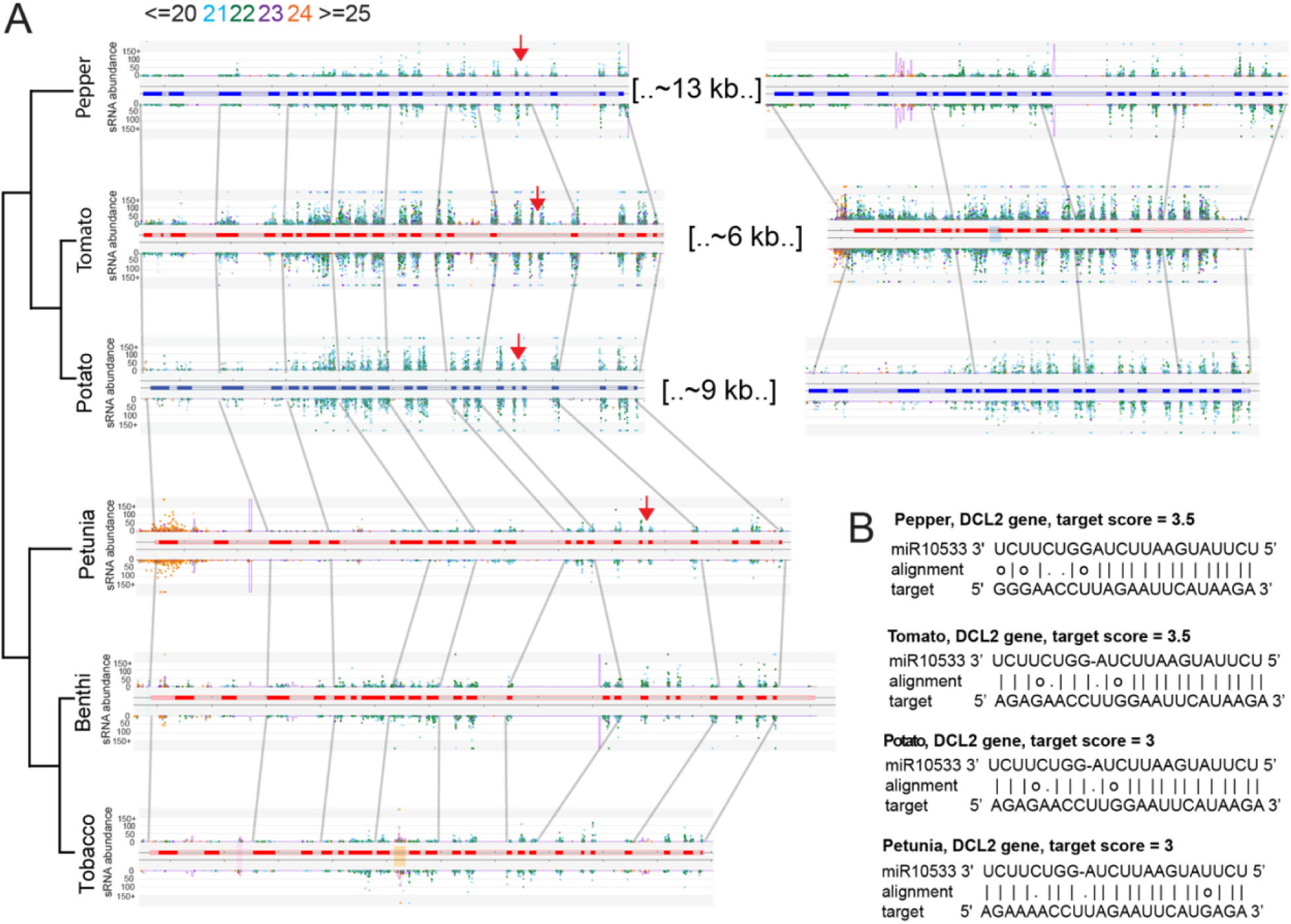
*DCL2* phasiRNA-producing loci are conserved across Solanaceous species. *DCL2* paralogs in pepper, tomato, and potato, and orthologs in petunia, benthi, and tobacco all produce 21-nt phasiRNAs (A). We identified conserved miRNA target sites for miR10533 and miR6026 in pepper, tomato, potato, and petunia (red and yellow arrow respectively denotes cleavage sites) that has a strong target score, and may be a conserved mechanism for triggering DCL2 *PHAS* locus production in these species (B).

### hc-siRNAs and transposable elements across SOL genomes

Transposable elements (TEs) are recognized as one of the major evolutionary forces that can shape genomes, via mutations and genome rearrangements. Presumably as protection from the possible detrimental consequences generated by TEs, plants have evolved a mechanism using 24-nt heterochromatic siRNAs (hc-siRNAs) that methylate and repress TEs (Ahmed et al., 2011; Wang et al., 2013). In order to analyze the hc-siRNA landscape in Solanaceae, we first identified TEs in all the available genomes. We observed that TEs represent between 25% (in petunia) and 48% of the genome (in benthi) (Supplemental Figure 5). As expected, the majority of TEs belong to an abundant LTR retrotransposon family, with most LTRs mapping specifically to Gypsy/Ty3 and Copia/Ty1 elements (Galindo-González et al., 2017) (Supplemental Figure 5A). Previous work mapped an expansion of LTRs in the Pepper genome; our results indicate that this observation applies broadly across the Solanaceae, and is likely a generalized signature of Solanaceous genomes (Park et al., 2012; Vicient and Casacuberta, 2017). After identifying TEs we analyzed the number of 24-nt reads that originate from each of the transposon superfamilies. Due to the repetitive nature of TEs, we did not attempt to further classify the sRNA reads into TE subfamilies. We observed that most of the sRNA reads mapped to retrotransposons, including LTR and non-LTR TEs (Supplemental Figure 5B). However, we were surprised to find that a large portion of hc-siRNAs also mapped to SINE elements. SINE elements comprise a unique subset of TEs; they are transcribed by RNA polymerase III, do not encode proteins, and are widespread in almost all multicellular eukaryotic genomes, with the exception Drosophila (Kramerov and Vassetzky, 2011). Our discovery of SINE-derived hc-siRNAs indicates that similar to LTRs, the spread of non-autonomous SINE elements is repressed via hc-siRNA-directed methylation.

Next, we analyzed the distribution of hc-siRNAs across Solanaceous genomes. We focused this analysis on tomato, pepper, potato and tobacco, which all have chromosome-scale assemblies. As expected, we found relatively high accumulation of hc-siRNAs along the telomeric ends of chromosomes; these hc-siRNAs promote telomere methylation in an RdDM–dependent manner (Vrbsky et al., 2010). We also observed a striking lack of hc-siRNAs across the entire potato genome (Figure 7). This result is similar to a previous observation described in strawberry, where, as in potato, asexual cloning is the primary means of propagation. Clonally propagated strawberry accumulates very low levels of DNA methylation relative to sexually propagated plants (Niederhuth et al., 2016). In the same way, we postulate that depleted hc-siRNA levels in potato could be caused by repeated rounds of asexual propagation. Further studies comparing sexually versus asexually propagated potato varieties would help resolve the connection between mode of reproduction and whether circumventing meiosis over multiple generations leads to degeneration of RNA-directed DNA methylation.

**Figure 7:**
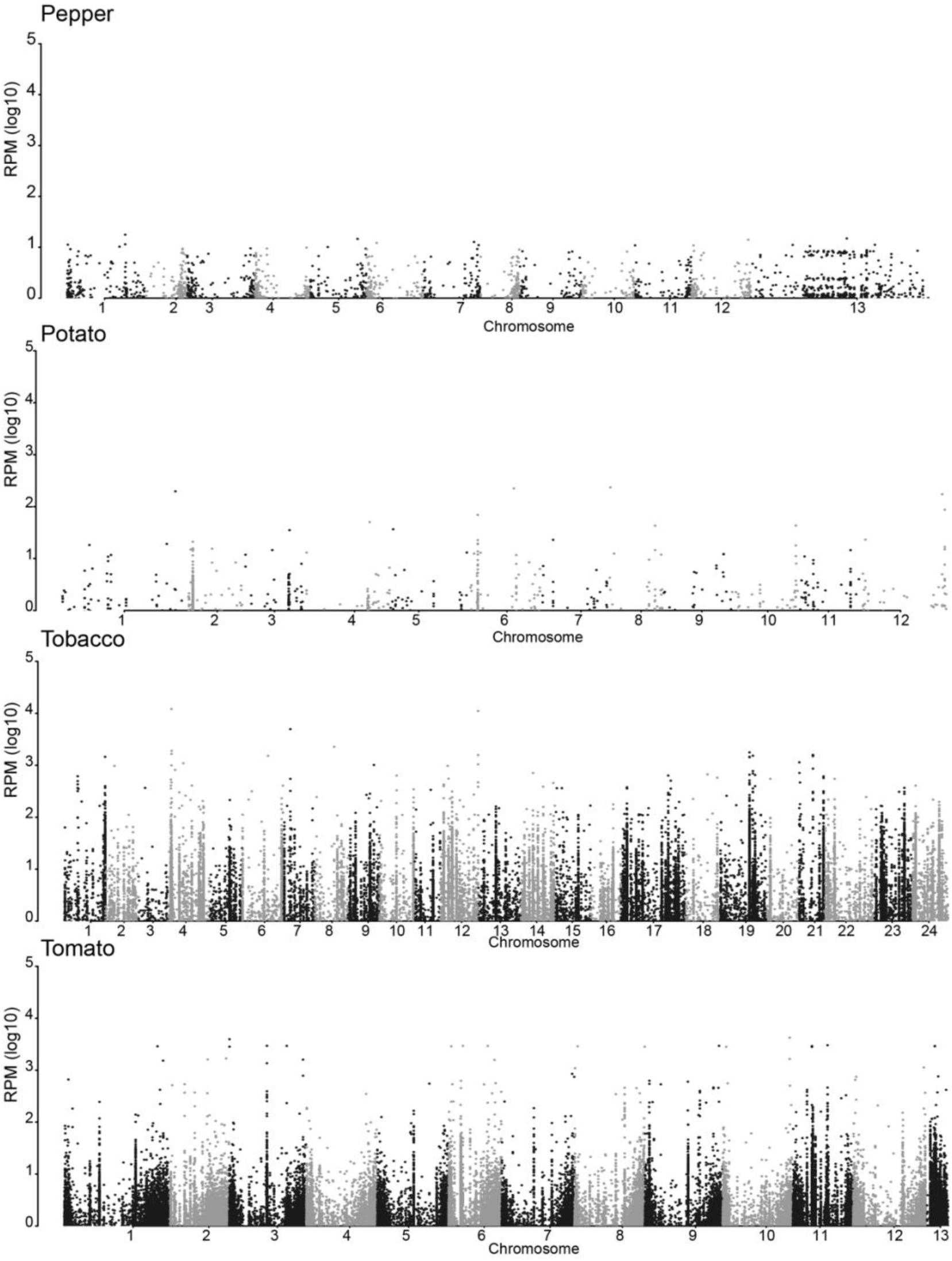
hc-siRNA expression across chromosomes in Solanaceae. Normalized read counts for 24-nt TE-associated hc-siRNAs are plotted across the assembled chromosomes for tobacco, tomato, potato, and pepper. The x-axis represents the chromosome number, alternating between grey and black, and the y-axis represents the average number of reads at each chromosome position. The average read number was calculated based on the number of available libraries for each species. As expected, the majority of hc-siRNAs map to telomeric ends. Unexpectedly, potato accumulates a low proportion of 24-nt hc-siRNAs relative to total sRNAs.

### Discovery of 22 nucleotide clusters expressed in response to geminivirus infection

A recent study in Arabidopsis characterized a class of 22-nt stress-responsive siRNAs that impact global translational repression (Wu et al. 2020). To test whether a similar class of sRNAs is present in the Solanaceae, we performed an analysis on our tomato expression libraries which contain the most extensive sampling with regard to abiotic and biotic stress treatments. We found a strongly-expressed cluster of 22-nt sRNA reads that mapped to a 4.8 kb region of the *S. lycopersicum* genome. Interestingly, this 22-nt sRNA cluster was uniquely expressed in a set of geminivirus infected samples (ToLCV-infected and TYLCV-infected), and absent from the rest of the tomato expression libraries, including mock-treated controls (Figure 8). This cluster of 22-nt sRNAs contains a *DISEASE RESISTANCE PROTEIN* (Solyc05g005330.3.1) and *ABSCISIC ALDEHYDE OXIDASE 3* (Solyc05g005333.1.1). To investigate whether there are additional 22-nt clusters that are specifically expressed in response to viral infection, we applied an intensive, genome-wide analysis to uncover highly expressed 22-nt sRNA clusters in the two viral-infected samples, using our mock treated libraries as controls. This genome-wide search uncovered 1,521 22-nt sRNA clusters in total, 390 of which passed our filtering criteria (Supplemental Figure 6 and Supplemental Dataset 10), of which, 22 showed more than a 4-fold induction in both viral treatments (Supplemental Figure 6B and Supplemental Dataset 10). Apart from the cluster shown in Figure 8A & B, one prominent example is the two clusters shown in Figure 8C & D, which are induced by 1,000-fold in response to ToLCV infection and by 10,000-fold in response to TYLCV. These highly-expressed, viral-induced clusters overlap with three gene copies of *EIN3-BINDING F-BOX PROTEIN 1* (Solyc05g008700.2.1, Solyc05g008720.1.1, and Solyc05g008730.1.1), one *MYZUS PERSICAE-INDUCED LIPASE 1* (Solyc05g008710.1.1), and one *RNA-DEPENDENT RNA POLYMERASE* (Solyc05g008740.1.1). Further studies are needed to test whether these geminivirus-induced 22-nt sRNA clusters have a functional role in plant defense to viral pathogens.

**Figure 8:**
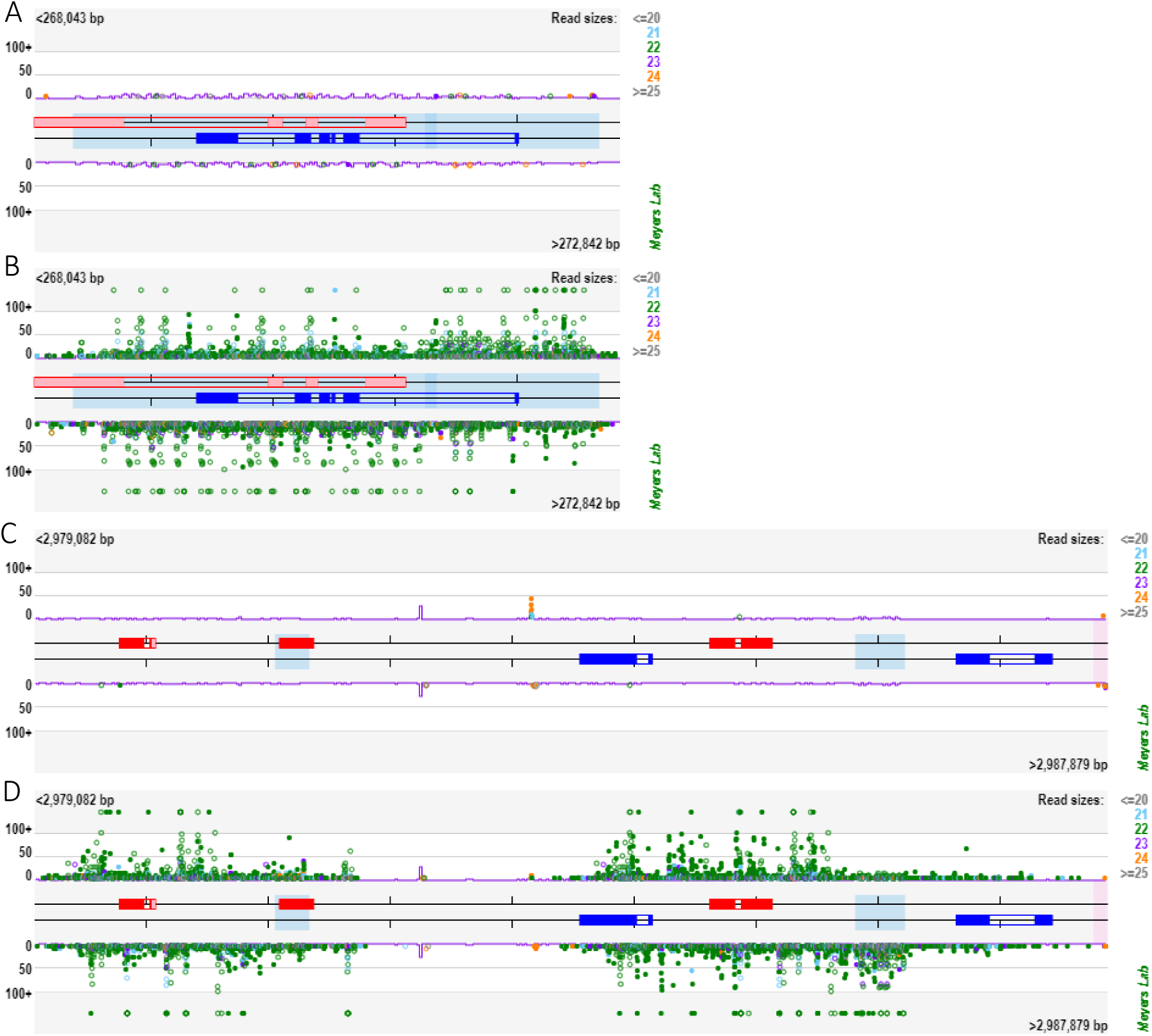
Tomato produces 22-nt sRNA clusters in response to geminivirus infection. Small RNA reads mapped to two tomato loci (Ch05:268043..272842 (A and B) and Ch05:2979082..2987879 (C and D)) for a gemnivirus infected and control sample pair shows strong expression of 22-nt siRNAs (shown as green dots) in response to viral treatments. The central red and blue boxes represent gene models from the 3.0 version of *S. lycopersicum* genome, and shaded boxes represent repetitive sequences. Dots represent sRNA reads, color-coded by length, mapped to the genomic region, with abundance represented by the position along the Y axis. Hollow dots are multi-mappers, and solid dots are unique-mappers. (A) and (C) are expression plots of control, mock-infected samples, and (B) and (D) are from the corresponding virus-infected samples.

## DISCUSSION

The Solanaceae family is widely used as a model for studying the evolutionary diversification of specialized metabolism (Fan et al., 2019), morphology (Koenig and Sinha, 2007; Martinez et al., 2020), fruit development (Klee and Giovannoni, 2011), and biotic and abiotic stress tolerance. In this study, we took advantage of the numerous publicly-available sRNA datasets that have been generated for the family to explore unique aspects of sRNA biology within the Solanaceae. We found several features that are shared amongst diverse Solanaceous species including a Solanales-wide duplication of sRNA biogenesis machinery, a previously-annotated Solanaceae-specific miRNA family that we show is actually a MITE-derived miRNA-like siRNA, and DCL2-derived 21-nt phasiRNAs that are broadly expressed across the diverse taxa in our study. We also found several phenomena that evolved within sub-family lineages of the Solanaceae. We discovered a *Nicotiana*-specific expansion of novel miRNAs, a distinct split in the number and types of phasiRNAs that are produced in the *Capsicum/Solanum* versus the *Petunia/Nicotiana* branches of the family, a dramatic depletion of 24-nt hc-siRNAs in asexually propagated potato, and 22-nt siRNA clusters that accumulate in response to viral infection in tomato. Below, we discuss some of the evolutionary implications for Solanaceae-wide and sub-family specific sRNA phenomena that we discovered in this study.

### Expansion and subfunctionalization of sRNA biogenesis machinery within the Solanaceae

In our reconstruction of sRNA biogenesis machinery machinery gene trees, we characterized lineage-specific expansions of *AGO1*, *AGO2* and *DCL2* gene families. While *AGO1* increased in copy number across the Solanaceae, *AGO2* and *DCL2* specifically expanded in the *Solanum*, and *Solanum/Capsicum* branches of the family, respectively, through tandem duplications. The tandem *DCL2* paralogs have been previously characterized to function as part of the tomato viral defense response (<u>Wang et al., 2018</u>), discussed in further detail below. Using public expression data for tomato, we identified distinct expression patterns in vegetative and reproductive organs for *DCL2* paralogs, with enriched abundance in reproductive organs, suggesting that these gene paralogs are undergoing subfunctionalization. Recent studies show that 24-nt phasiRNAs are produced only in reproductive organs of angiosperms, while 21-nt phasiRNAs are produced in both vegetative and reproductive tissues (Xia et al, 2019, Pokhrel et al., 2020). The 24-nt reproductive phasiRNAs are triggered by miR2275 in most of the angiosperms while in Solanaceous species, the miRNA trigger remains unknown (Xia et al, 2019, Pokhrel et al, 2021). Given that *DCL2* was shown to process endogenous 22-nt miRNAs, and that the expression of some *DCL2* paralogs peak in reproductive organs, it would be interesting to functionally test whether this *DCL2* paralog expressed in reproductive tissues specifically processes a 22-nt miRNA that triggers reproductive 24-nt phasiRNA biogenesis.

### Solanaceae-wide miR5303 cluster is derived from MITEs

Our cross-species miRNA predictions and clustering revealed 12 family-wide clusters of miRNAs. While the majority of these clusters are associated with deeply-conserved land plant miRNA families that are shared outside of the Solanaceae, we investigated the Solanaceae-specific cluster annotated as miR5303 (Mohorianu et al., 2011). We discovered that this family exhibits many features that are atypical of miRNAs. For example, related miR5303 sequences in miRBase include both 21-nt and 24-nt mature sequences and most of the Solanaceae genomes harbor relatively high copy numbers of miR5303 isomiRs that putatively target thousands of genes based on our miRNA-target prediction analysis. We found that, similar to a previous study on MITE-derived miRNA-like siRNAs, mature miR5303 “miRNA” sequences align perfectly to the terminal inverted repeat (TIR) region of Solanaceae MITEs. (Kuang et al., 2009). How did our miRNA prediction pipeline lead to the identification of MITE-derived sRNAs? MITEs have near perfect hairpin folding and express sRNAs within mature miRNA size ranges, and thus fit many of the criteria used to predict novel miRNA precursors. It has been proposed that MITEs may contribute to the *de novo* evolution of miRNAs. However, this model is somewhat controversial, as MITEs tend to express imprecise sRNAs that are variable in both length and sequence, and thus are more likely the products of siRNA rather than miRNA biogenesis machinery (Cui et al, 2017). Even so, the conspicuous presence of miR5303-related sRNAs exclusively in the Solanaceae is intriguing, and further investigation into putative functional roles for these siRNAs could be interesting.

### *DCL2* is a broadly expressed *PHAS* locus that may control viral responses across the Solanaceae

In a previous study focused on mechanisms of viral resistance in tomato, paralogous copies of *DCL2* were shown to process a 22-nt miRNA (miR6026) that has dual functions, participating in antiviral responses and creating a self-regulating feedback loop by triggering *DCL2* phasiRNA production (<u>Wang et al., 2018</u>). In this study we identified *DCL2* as a widely-expressed *PHAS* locus that is present in all seven Solanaceous species in our study. Contrary to previous characterization of this locus, we predicted that another 22-nt miRNA, miR10533, triggers phasiRNA production in *Solanum*, *Capsicum*, and *Petunia* genera. Similar to miR6026, miR10533 targets disease resistance-related genes in addition to *DCL2* (Supplemental Dataset 8). Based on the accumulation patterns of *DCL2* phasiRNAs, neither miRNA appears to be an obvious trigger, although target mimic experiments with miR6026 showed reduced *DCL2* phasiRNA production (Wang et al., 2018). We identified the miR6026 site more than 250 bp outside of our predicted phasiRNA production region, whereas miR10533 is sandwiched within the phasiRNA register (Figure 6). The lack of an obvious trigger suggests two possibilities: 1) there may be another, as yet unidentified miRNA trigger, and/or 2) as hypothesized by previous authors, the 22-nt sRNA products generated from the *DCL2* paralogs may be cross triggering one another in *cis*, creating multiple phasiRNA initiation sites (Wang et al., 2018). We have performed extensive miRNA predictions for tomato, given the depth of sRNA sequence data available for this species, and thus it is unlikely that we are simply missing an additional trigger for *DCL2* phasiRNA production. Consistent with the second hypothesis, we observed weaker phasing scores across most of the *DCL2 PHAS* loci, indicating that cross-locus regulatory interactions trigger phasiRNA production across *DCL2* loci.

*DCL2 PHAS* loci have also been identified in two different Fabaceae models, soybean and medicago, where they are triggered by independent miRNAs (Li et al., 2010; Zhai et al., 2011). Our study, in combination with this previous work, shows that phasiRNA target-trigger interactions around *DCL2* have evolved multiple times within and across eudicot families. This evolutionary hotspot for phasiRNA production fits into a model in which the expression of sRNA biogenesis machinery is regulated via self-triggered phasiRNA production. Depending on the species, different components of sRNA biogenesis machinery are processed into *PHAS* loci. In soybean, for example, *SGS3* is targeted for phasiRNA production (Song et al., 2011), and in Arabidopsis, *AGO1* and *DCL1* are both processed into *PHAS* loci (Xie et al., 2003; Vaucheret et al., 2004). Why do plants use post-transcriptional regulation to control siRNA biogenesis machinery? One potential theory is that by regulating sRNA biogenesis machinery at the post-transcriptional level, plants can maintain a standing pool of transcripts for sRNA biogenesis that can be rapidly released and translated in response to pathogens.

### Lineage-specific expression of miRNAs and phasiRNAs

One of our goals in this study was to identify miRNAs that are uniquely conserved within the Solanaceae; however, as discussed above, we demonstrate that miR5303, the only Solanaceae-specific “miRNA family” that we were able to predict in all of our study species, is actually a MITE-derived siRNA. Rather than uncovering broadly conserved miRNAs, we discovered numerous instances of lineage-specific miRNAs, with the most pronounced example coming from our representative species for *Nicotiana*. Even after stringent filtering, we predicted over 350 miRNAs in the benthi genome, with more than half of these comprising previously unannotated, lineage-specific miRNAs. Congruent with previous observations, we show that these lineage-specific miRNAs are generally expressed at lower levels than deeply conserved family members (Supplemental Figure 2) (Axtell et al., 2007). There are, however, a handful of newly annotated miRNAs that are abundantly expressed (>1,000 rpm; Supplemental Figure 2). Given the ease of agrobacterium infiltration in benthi, followup studies using transiently expressed target mimic knockdowns and PARE sequencing may provide new insight into the targets and putative functions of these rapidly-evolving *Nicotiana*-specific miRNAs. One bottleneck that we encountered in this analysis was a lack of deep sRNA sequence data for tobacco. Despite the lack of sampling in tobacco, we still predicted more than 150 miRNAs for this genome, which is similar to the number of miRNAs that we found in the tomato genome, despite having almost 10 times more sRNA sequence data for tomato. We are likely looking at the tip of the iceberg when it comes to miRNA discoveries in tobacco, and we predict that deeper sequencing in tobacco and related *Nicotiana* species will facilitate big gains when it comes to miRNA discovery in the Solanaceae.

In a reverse trend, our genome-wide 21-nt phasiRNA predictions uncovered relatively few *PHAS* loci for *Nicotiana* and *Petunia* species compared to the *Solanum/Capsicum* branch of the family (Figure 5A). A distinct lack of *PHAS* loci encoding NB-LRRs in *Nicotiana* and *Petunia* accounts for one of the biggest differences in the number of *PHAS* loci between these two branches of the family. We also discovered two strongly expressed non-coding *PHAS* loci that are conserved at the sub-family level. One is conserved in both benthi and tobacco, and was previously described to function in the post-transcriptional regulation of PPR genes (Supplemental Figure 4) (Xia et al., 2013). The other locus is uncharacterized, and specific to tomato and potato (Figure 5B). We discovered that this new *PHAS* locus produces a prominent 21-nt siRNA that putatively targets MYB transcription factors (Figure 5D). It would be interesting to test whether this Solanum-specific *PHAS* locus really does function in regulating the expression of MYB transcription factors. PhasiRNA production is triggered by miRNAs. Therefore, it is logical that in addition to discovering lineage-specific miRNAs, we also found lineage-specific *PHAS* loci across the Solanaceae.

### Repressed hc-siRNA expression reflects a history of asexual propagation in potato

Our genome-wide hc-siRNA analysis revealed a low abundance of 24-nt TE-associated siRNAs in potato. We hypothesize that this relative depletion in potato is caused by the asexual means of propagation that is common for this crop. Epigenetic marks are faithfully copied during mitosis; however, during meiosis, they are reset by RdDM-based repatterning, which involves TE-derived hc-siRNAs directing DNA methylation (Daxinger et al., 2009; Schoft et al., 2009; Mosher and Melnyk, 2010; Slotkin et al., 2009; Verhoeven and Preite, 2014). Asexual propagation circumvents meiosis, and thus this RdDM-based reseting of epigenetic marks is avoided (Verhoeven and Preite, 2014), and over many rounds of asexual propagation DNA methyl-cytosine marks and their associated hc-siRNAs may become depleted. In line with this hypothesis, a related study investigating genome-wide DNA methylation patterns in diverse species found a stark depletion in CHH cytosine methylation in strawberry, another asexually propagated species (Niederhuth et al., 2016). However, there are limitations to our conclusions regarding depleted hc-siRNAs in potato. These limitations include imperfect sampling, notably, the majority of our potato samples are from vegetative tissues (although this is true for most of the species in this study; Supplemental Dataset 1), and insufficient information about the source tissue that was used to generate these public datasets, particularly information regarding how many generations of asexual reproduction the source tissue underwent prior to sampling. Since sRNA-derived TE silencing gradually accumulates over multiple generations, a well-devised study comparing long-term asexually versus sexually propagated source tissue would help resolve whether levels of RdDM are directly impacted by assexual propagation, and whether these hc-siRNAs could be reactivated during reproductive development.

### 22-nt clusters expressed in response to viral infection in tomato

Recent work in Arabidopsis uncovered 22-nt siRNAs that are expressed in response to environmental stress, and are capable of impacting global translational repression, including the translation of their cognate genes. Moreover, this translational repression is correlated with an inhibition of growth and activation of stress-response pathways (Wu et al. 2020). In this study, we identified similar 22-nt siRNA clusters in tomato that are expressed in response to viral infection. Whether these siRNAs are also involved in attenuating global translation and growth so that more resources can be allocated to antiviral response remains to be seen. Several of the clusters that we identified are generated from genes involved in ABA production and ethylene signaling, indicating that these 22-nt siRNA clusters may coordinate genomic responses to pathogen attack by fine tuning the expression of ABA and ethylene pathways. Future work investigating whether this response is initiated by the plant in order to activate defense mechanisms or perhaps is hijacked by the virus in order to silence antiviral responses would provide insight into the evolutionary function of these distinctive siRNAs.

## CONCLUSION

In conclusion, while genera within the Solanaceae share unifying features in the evolutionary history of their sRNA biology, there are many lineage-specific phenomena that are not broadly shared across the family. This conclusion is important as it provides valuable information on the appropriateness of a given model species for providing generalized insights into sRNA biology. Simultaneously, we encountered limitations due to disparate sampling within the Solanaceae. For example, while sample data exist across developmental stages, environments, and genotypes to support investigations into tomato sRNA biology, there is a lack of publicly-available resources to study tomato’s sister species, eggplant (Chapman, 2019). Similarly, our understanding of sRNA biology for *Petunia hybrida* is largely limited to reproductive samples. Given the sub-family phenomena that we identified in this study, we suggest that delving into under-sampled taxa rather than increasing sampling in well-studied species will yield a greater return on investment when it comes to discovering new aspects of sRNA biology within the Solanaceae.

## MATERIALS AND METHODS

### Evolution of small RNA biogenesis protein genes in the Solanaceae

We downloaded protein sequence files from Phytozome v13, the Sweetpotato Genomics Resource, and the SOL Genomics Network databases, and selected proteomes for *Arabidopsis thaliana*, two Convolvulaceae (*Ipomoea trifida* and *Ipomoea triloba*) and the seven Solanaceae species (*Nicotiana benthamiana*, *Nicotiana tabacum*, *Petunia axillaris*, *Petunia inflata*, *Capsicum annuum*, *Solanum lycopersicum* and *Solanum tuberosum*) for analysis (Supplemental Table 2). We used SonicParanoid v1.2.6 (<u>Cosentino and Iwasaki, 2018</u>) and OrthoFinder v2.3.11 (<u>Emms and Kelly, 2015</u>) to perform a gene orthology inference using default parameters. To identify orthologous groups related to NRP, RDR, DCL, DRB and AGO proteins, we used the reference protein IDs available in the UniProt database for *A. thaliana*. We then aligned the retrieved protein sequences using default parameters in MUSCLE v3.8.1551 (<u>Edgar, 2004</u>), and trimmed alignments with trimAL v1.4.rev15 applying the -gappyout option (<u>Capella-Gutierrez et al., 2009</u>). To infer phylogenetic trees we implemented the maximum likelihood method in IQ-TREE v1.6.12 with the following parameters: -alrt 1000 and -bb 1000 (<u>Minh et al., 2013</u>; <u>Nguyen et al., 2015</u>; <u>Kalyaanamoorthy et al., 2017</u>). We used iTOL v4 (<u>Letunic and Bork, 2019</u>) to visualize consensus trees and summarize orthologous protein genes for the five gene families investigated in each species (Supplemental Table 1). To investigate organ-specific expression for the DCL1, AGO1, and AGO2 paralogs identified for tomato, we downloaded expression data for *Solanum lycopersicum* from The Bio-Analytic Resource for Plant Biology (<u>Waese et al., 2017</u>).

### Data collection, quality filters, and read processing

We downloaded raw sRNA seq and PARE libraries for *Solanum lycopersicum* (tomato), *S. tuberosum* (potato), *Nicotiana benthamiana*, *N. tabaccum* (tobacco), *Capsicum annum* (pepper), *Petunia axillaris/P. inflata* (petunia) from the NCBI SRA (https://www.ncbi.nlm.nih.gov/sra; detailed information regarding raw and processed libraries can be found in Supplemental Dataset 1. Briefly, we processed raw FASTQ reads using a Python script that performs quality filtering, trimming, adapter removal, and then converts reads into chopped tag count files (https://github.com/atulkakrana/preprocess.seq (Bolger et al., 2014)). We then mapped our reads to their respective reference genomes (Supplemental Table 2) using Bowtie 1 (Langmead, 2010) allowing for zero mismatches. We applied two criteria for filtering out low-quality data: first, we excluded libraries with low mapping rates, and second, we removed libraries that lacked a significant accumulation of 21-nt and 24-nt reads based on size distribution plots (Supplemental Figure 7), indicating potential RNA degradation. We assembled species-specific expression databases using these final, high-quality, processed datasets (Nakano et al., 2020).

### Solanaceae genome versions and functional gene annotations

We used the following reference genomes and their associated annotation files to build expression databases and perform the bioinformatic analyses described in this study: *Capsicum annum* Zunla-1 v2.0 (http://public.genomics.org.cn/BGI/pepper/) (Qin et al., 2014), *Petunia axillaris* and *P. inflata* (Bombarely et al., 2016), *Nicotiana tabacum* v4 (Edwards et al., 2017), *Nicotiana benthamiana* v 0.3 (Bombarely et al., 2012), *Solanum tuberosum* (Potato Genome Sequencing Consortium, 2011), and *Solanum lycopersicum* build 2.5 (Tomato Genome Consortium, 2012) (ftp://ftp.solgenomics.net/genomes/).

### MicroRNA Predictions

We used three different software packages to obtain a robust prediction of microRNAs (miRNAs) across our Solanaceous model genomes: ShortStack (Axtell, 2013), miR-PREFeR (Lei and Sun, 2014), and miRPD (a version of miReap optimized for plant miRNA prediction; (Sun et al., 2013)). We standardized the minimum mapping coverage and size of hits across the three programs in order to control for dissimilarities in software performance. The vast majority of miRNAs fall between 20-22 nt in length so we narrowed our predictions to this 20-22 nt size range. Each program employs a different method for calculating coverage. In ShortStack we set the minimum coverage to 15, while in miR-PREFeR and miRPD we set the minimum coverage to 20. We then filtered out predicted miRNA precursors based on recently published guidelines for miRNA predictions (Axtell and Meyers, 2018). Briefly, we required that all miRNAs express both strands of the miRNA duplex (the mature miRNA and miRNA-star), and originate from a precursor that folds with a normalized minimum free energy (MFE) of < -0.2 kcal/mol/nucleotide. Notably, enforcing the expression of a complementary miRNA-star sequence resulted in little to no filtering, so we applied a more stringent filter requiring that the abundance of the complementary miRNA-star reads were ≥10% of the abundance of the mature miRNA.

We applied additional filtering for “novel” miRNAs that did not match miRBase entries (see “miRNA conservation across species,” below). To remove hairpin structures that might originate from MITEs and tRNAs, we filtered out novel miRNAs that aligned to our list of annotated transposable elements (TEs) and tRNA datasets for the Solanaceae (Supplemental Dataset 11). We found a surprising number of novel miRNAs in our *Nicotiana benthamiana* (hereafter, “benthi”) dataset which warranted manual curation. To investigate whether our novel benthi miRNAs originate from canonical precursor structures, we mapped our shortread data onto predicted miRNA precursors using structVis v0.4 (github: https://github.com/MikeAxtell/strucVis), and removed novel miRNAs that showed non-canonical folding and/or miRNA/miRNA* expression. After these filtering steps we found the miRPD results to be highly inconsistent, producing unrealistically high numbers of predicted miRNAs for some of our species and zero predicted miRNAs for others. Thus, we decided to limit our analyses to the ShortStack and miR-PREFeR results. To create a consensus set of miRNAs for these two prediction programs, we collapsed precursors with overlapping coordinates from these two software programs into unique entries. Overall, we found that ShortStack was more conservative, predicting fewer miRNAs than miR-PREFeR; however, the miRNAs were also predicted with appreciable confidence and resulted in minimal post-prediction filtering. Our hybrid prediction approach in combination with stringent filtering allowed for sensitive, yet robust predictions across the species.

### MicroRNA conservation across species

To facilitate cross species comparisons of conserved and novel miRNAs within the Solanaceae, we first clustered mature miRNA sequences for the predicted miRNAs (described above) into families by requiring pairwise sequence overlap of ≥75%. To annotate the miRNA families, we aligned mature miRNA sequences to the latest v22 miRBase database of all land plants requiring an 85% match, and provided new IDs for novel miRNA families (Kozomara and Griffiths-Jones, 2014; Kozomara et al., 2019). We tabulated all novel and previously annotated miRNA families across the Solanaceae in a master table that reports presence and absence for each miRNA family (Supplemental Dataset 5).

### *PHAS* locus predictions and annotations

We used three programs to identify *PHAS* loci: ShortStack (Axtell, 2013), PHASIS (Kakrana et al., 2017), and PhaseTank (Guo et al., 2015). We ran PHASIS and PhaseTank using the default parameters listed in the program manuals and filtered the results for high-confidence loci that were identified with a p-value of ≤.001. We ran ShortStack using the same parameters that were applied to our miRNA predictions. PhaseTank did not perform consistently across the analyzed genomes; it only reported *PHAS* loci for unassembled regions of the genomes. After we removed unassembled contigs, the program returned no results, so we decided to remove PhaseTank from our analysis pipeline. We merged the *PHAS* loci obtained from PHASIS and ShortStack into a single list for each species, and collapsed loci located within 400 nucleotides of each other into individual *PHAS* predictions (Supplemental Dataset 7).

We annotated coding versus non-coding *PHAS* loci using available CDS GFF files. To characterize non-coding *PHAS* loci we aligned our non-coding *PHAS* loci to the NCBI non-redundant (nr) database and annotated the loci with one of the top ten hits using default alignment parameters in BLAST (Altschul et al., 1990). To test whether coding *PHAS* loci were generated from orthologous genes across species, we constructed gene orthogroups for species with associated peptide fasta files. Finally, we linked these orthogroup predictions with annotation files for tomato, Arabidopsis, and pepper (Supplemental Dataset 9). The published pepper gff file lacks annotations, so we generated our own pepper annotation file by running a BLASTP search against the NCBI NR database and retrieving the best BLAST hit for each entry.

### miRNA target predictions

We used sPARTA, a miRNA target prediction program to identify miRNA targets with the following specifications: -genomeFeature 0 -tarPred H -tarScore S (Kakrana et al., 2015). We applied a cut off penalty score of ≤4 to filter for reliable miRNA-mRNA target interactions in benthi (Supplemental Dataset 4) and tomato (Supplemental Dataset 6). *PHAS* loci are triggered by 22 bp miRNAs that have a 5’ Uracil bias (Chen et al., 2010; Cuperus et al., 2010). We ran sPARTA with the -featureFile option to identify miRNAs from our predicted miRNA list that trigger *PHAS* locus production from our predicted PhasiRNA list. We applied a penalty score of ≤4 to filter for PHAS locus triggers, prioritized miRNA triggers that start with Uracil (U), and selected miRNA triggers equal to 22 bp with the exception of miR390 (Supplemental Dataset 8).

### Hc-siRNAs and transposable elements across SOL genomes

To analyze hc-siRNAs, we first ran RepeatMasker (http://www.repeatmasker.org) on each one of the analyzed genomes using the Viridiplantae RepBase database (http://www.girinst.org/repbase) as a library, to identify transposable elements (TE). Next, we used the identified TEs as a database to map sRNA reads using Bowtie (Langmead, 2010) with default parameters and categorized all 24-nt long reads that mapped to any of our identified TEs as hc-siRNAs. It was impossible to identify the specific origin of these sRNAs due to the repetitive nature of TEs, so we only considered TE superfamilies for this study. Genome-wide representation of hc-siRNAs was visualized by plotting Manhattan plots with the qqman R package (Turner, 2014). We plotted the data based on the genome mapping coordinate along the chromosome for each hc-siRNA and its relative abundance.

### Identification of 22-nt clusters expressed in response to geminivirus infection

To discover 22-nt sRNA clusters that may be linked to geminivirus infection in tomato, we downloaded sRNA libraries for control and infected treatments from the TOMATO_sRNA_SOL Next-Gen DB (Nakako et al., 2020) and filtered for 22-nt reads. We tuned the parameters for 22-nt cluster identification using two fruit peel libraries (Tomato_S_7 and Tomato_S_8, cv. Ailsa Craig, Gao et al., 2015) and two leaf libraries (Tomato_S_50, cv. Moneymaker, and Tomato_S_53, line FL505, Bai et al., 2016). To identify clusters that accumulate in response to geminivirus infection, we analyzed two infected leaf libraries (Sly_leaves_t2, cv. Pusa Ruby, Saraf et al., 2015; Tomato_S_52, cv. Moneymaker, Bai et al., 2016) and their corresponding control libraries (Sly_leaves_t1 and Tomato_S_50, respectively). We applied the following analysis pipeline to identify 22-nt sRNA clusters: first, we ran ShortStack with the parameters tuned to yield the highest reproducibility between repeated runs, relatively high similarity between results from similar samples (e.g. leaf vs. leaf), and relatively high difference between results from distinct samples (e.g. leaf vs. fruit peel), without compromising the alignment rate (Johnson et al., 2016). This fine tuning resulted in the following parameter set: --mmap u –mismatch 1 –bowtie_m 3 –ranmax 2 –pad 125 –mincov 75rpm. Using these parameters, we ran ShortStack to identify all of the 22-nt sRNA clusters in the aforementioned four libraries. Next, we filtered out clusters that were shorter than 200 bp in length, and clusters that were inconsistently expressed in one infected-control sample pair, but not in the other. We applied a threshold of absolute log2 fold change greater than two in order to identify differentially expressed clusters between the geminivirus infected and mock-treated samples.

### Data visualization and representation

We visualized the 22-nt sRNA clusters using the transcriptome browser function in the TOMATO_sRNA_SOL Expression database (Nakano et al., 2006; Nakano et al., 2020; https://mpss.meyerslab.org/). We generated dot plots using the ggplot2 package (Wickham, 2016), heatmaps using gplots package (Warnes et al., 2019), and heatmaps of DCL2 and AGO1 paralog expression using the TBTools heatmap function (Chen et al., 2018a).

## AUTHOR CONTRIBUTIONS

B.C.M and M.H.F. conceived the original research plan and supervised the analyses, and supervised and completed the writing; P.B., S.B., S.K., S.P., S.T., C.T., C.S., H.R., S.G.R.G, P. G. and P.P. performed the analyses, generated tables and figures, and contributed to writing of the article; M.N and A.D. provided technical assistance with data processing. M.H.F. agrees to serve as the author responsible for contact and ensures communication.

## SUPPLEMENTAL DATA

Supplemental Dataset 1 – SRA Sample Processing

Supplemental Dataset 2 – miRNAs Annotated by Species

Supplemental Dataset 3 – Benthi structVis Filtering

Supplemental Dataset 4 – Benthi miRNA Target Predictions

Supplemental Dataset 5 – Solanaceae miRNA Families

Supplemental Dataset 6 – Tomato miRNA Target Predictions

Supplemental Dataset 7 – phasiRNAs Annotated by Species

Supplemental Dataset 8 – PHAS Locus miRNA Interactions

Supplemental Dataset 9 – Solanaceae Orthogroups

Supplemental Dataset 10 – Table of 22-nt sRNA Clusters

Supplemental Dataset 11 – Solanaceae TEs

## SUPPLEMENTAL TABLES

**Supplemental Table 1:**
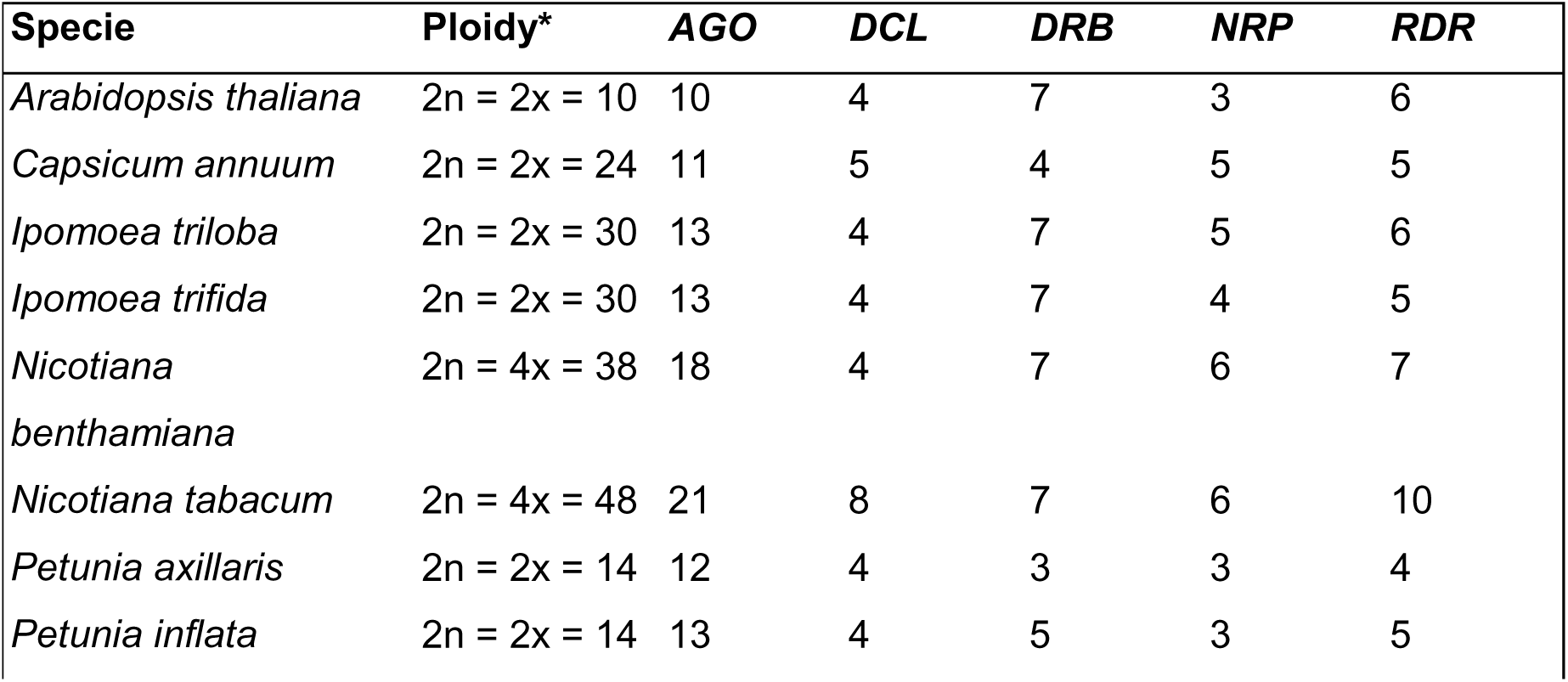
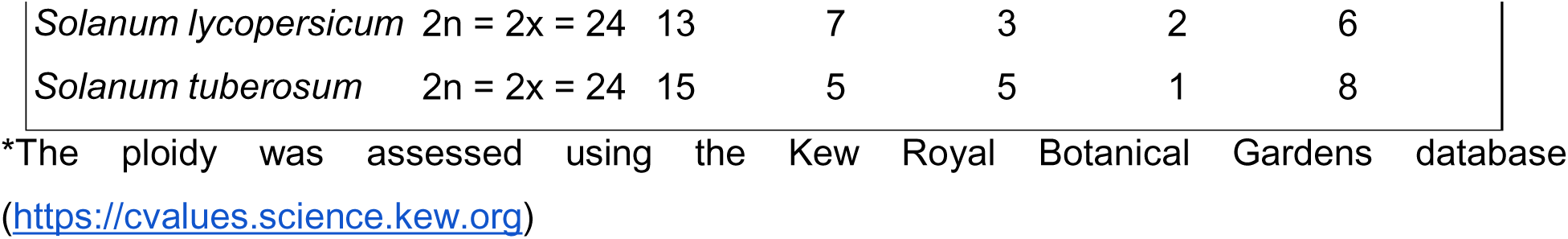
Summary of annotated *AGO*, *DCL*, *DRB*, *NRP* and *RDR* family members in Solanaceous and outgroup genomes.

| Specie | Ploidy* | *AGO* | *DCL* | *DRB* | *NRP* | *RDR* |
| --- | --- | --- | --- | --- | --- | --- |
| *Arabidopsis thaliana* | 2n = 2x = 10 | 10 | 4 | 7 | 3 | 6 |
| *Capsicum annuum* | 2n = 2x = 24 | 11 | 5 | 4 | 5 | 5 |
| *Ipomoea triloba* | 2n = 2x = 30 | 13 | 4 | 7 | 5 | 6 |
| *Ipomoea trifida* | 2n = 2x = 30 | 13 | 4 | 7 | 4 | 5 |
| *Nicotiana benthamiana* | 2n = 4x = 38 | 18 | 4 | 7 | 6 | 7 |
| *Nicotiana tabacum* | 2n = 4x = 48 | 21 | 8 | 7 | 6 | 10 |
| *Petunia axillaris* | 2n = 2x = 14 | 12 | 4 | 3 | 3 | 4 |
| *Petunia inflata* | 2n = 2x = 14 | 13 | 4 | 5 | 3 | 5 |
| *Solanum lycopersicum* | 2n = 2x = 24 | 13 | 7 | 3 | 2 | 6 |
| *Solanum tuberosum* | 2n = 2x = 24 | 15 | 5 | 5 | 1 | 8 |

\*The ploidy was assessed using the Kew Royal Botanical Gardens database (https://cvalues.science.kew.org)

**Supplemental Table 2:**
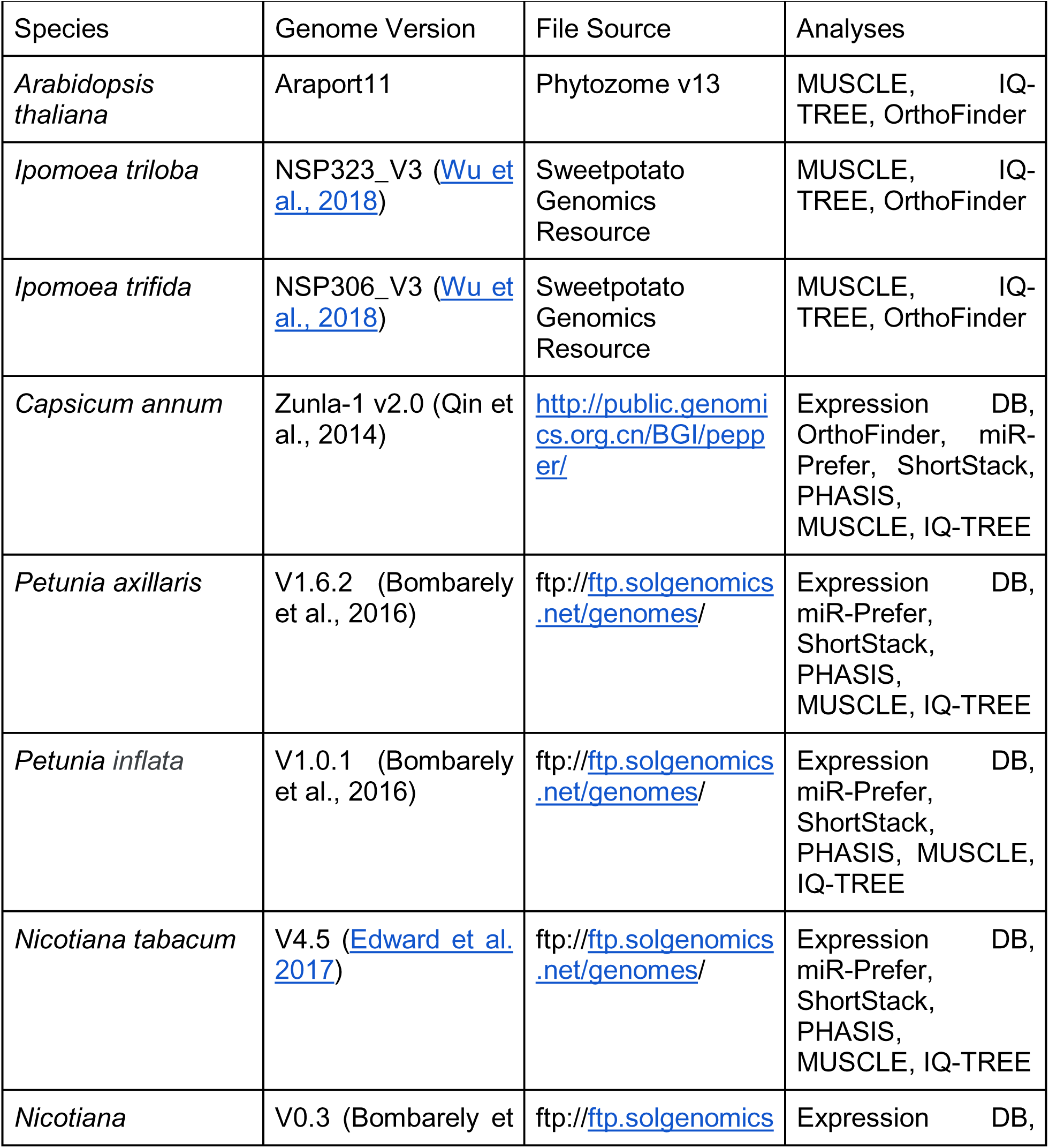
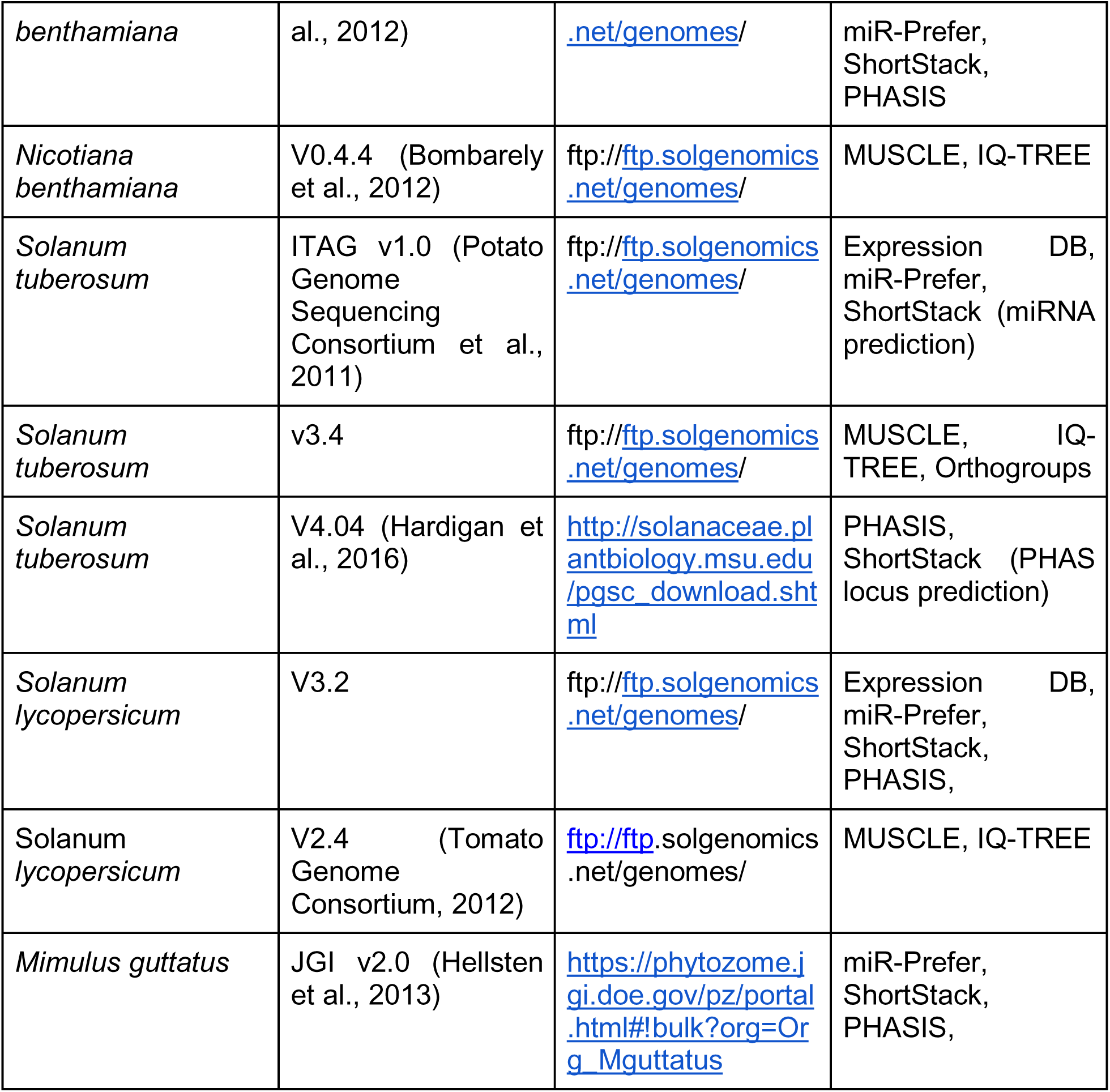
Reference genomes and software used for bioinformatic analyses.

## ACKNOWLEDGEMENTS

We thank Rui Xia for generating the annotation file that we used for the pepper (*Capscium annuum*) genome, and Feray Demirci and Deepti Ramachandruni for assistance with aspects of data processing. The work was supported with funding from NSF IOS awards 1842698, 754097 and 1650843, and USDA NIFA award 2019-67013-29010 to Meyers; NSF IOS awards 1523668 and 1942437 to M. Frank.

## REFERENCES

Ahmed I, Sarazin A, Bowler C, Colot V, Quesneville H (2011) Genome-wide evidence for local DNA methylation spreading from small RNA-targeted sequences in Arabidopsis. Nucleic Acids Res 39: 6919–6931

Altschul SF, Gish W, Miller W, Myers EW, Lipman DJ (1990) Basic local alignment search tool. J Mol Biol 215: 403–410

Axtell MJ (2018) The small RNAs of Physcomitrella patens : Expression, function and evolution. Annual Plant Reviews online 113–142

Axtell MJ (2013) ShortStack: comprehensive annotation and quantification of small RNA genes. RNA 19: 740–751

Axtell MJ, Meyers BC (2018) Revisiting Criteria for Plant MicroRNA Annotation in the Era of Big Data. The Plant Cell 30: 272–284

Axtell MJ, Snyder JA, Bartel DP (2007) Common functions for diverse small RNAs of land plants. Plant Cell 19: 1750–1769

Axtell MJ, Westholm JO, Lai EC (2011) Vive la différence: biogenesis and evolution of microRNAs in plants and animals. Genome Biol 12: 221

Bolger AM, Lohse M, Usadel B (2014) Trimmomatic: a flexible trimmer for Illumina sequence data. Bioinformatics 30: 2114–2120

Bombarely A, Moser M, Amrad A, Bao M, Bapaume L, Barry CS, Bliek M, Boersma MR, Borghi L, Bruggmann R, et al (2016) Insight into the evolution of the Solanaceae from the parental genomes of Petunia hybrida. Nat Plants 2: 16074

Bombarely A, Rosli HG, Vrebalov J, Moffett P, Mueller LA, Martin GB (2012) A draft genome sequence of Nicotiana benthamiana to enhance molecular plant-microbe biology research. Mol Plant Microbe Interact 25: 1523–1530

Canto-Pastor A, Santos BA, Valli AA, Summers W, Schornack S, Baulcombe DC Enhanced resistance to bacterial and oomycete pathogens by short tandem target mimic RNAs in tomato. doi: 10.1101/396564

Capella-Gutierrez S, Silla-Martinez JM, Gabaldon T (2009). trimAl: a tool for automated alignment trimming in large-scale phylogenetic analyses. Bioinformatics 25:1972–1973.

Cardoso TC de S, Alves TC, Caneschi CM, Santana DDRG, Fernandes-Brum CN, Reis GLD, Daude MM, Ribeiro THC, Gómez MMD, Lima AA, et al (2018) New insights into tomato microRNAs. Sci Rep 8: 16069

Chapman MA (2019) Introduction: The Importance of Eggplant. In MA Chapman, ed, The Eggplant Genome. Springer International Publishing, Cham, pp 1–10

Chen C, Chen H, He Y, Xia R (2018a) TBtools, a Toolkit for Biologists integrating various biological data handling tools with a user-friendly interface. bioRxiv 289660

Chen C, Zeng Z, Liu Z, Xia R (2018b) Small RNAs, emerging regulators critical for the development of horticultural traits. Horticulture Research 5: 6–8

Chen H-M, Chen L-T, Patel K, Li Y-H, Baulcombe DC, Wu S-H (2010) 22-Nucleotide RNAs trigger secondary siRNA biogenesis in plants. Proc Natl Acad Sci U S A 107: 15269–15274

Cosentino S, Iwasaki W (2018). SonicParanoid: fast, accurate and easy orthology inference. Bioinformatics. 35(1):149–51.

Cuperus JT, Carbonell A, Fahlgren N, Garcia-Ruiz H, Burke RT, Takeda A, Sullivan CM, Gilbert SD, Montgomery TA, Carrington JC (2010) Unique functionality of 22-nt miRNAs in triggering RDR6-dependent siRNA biogenesis from target transcripts in Arabidopsis. Nat Struct Mol Biol 17: 997–1003

Daxinger L, Kanno T, Bucher E, van der Winden J, Naumann U, Matzke AJM, Matzke M (2009) A stepwise pathway for biogenesis of 24-nt secondary siRNAs and spreading of DNA methylation. EMBO J 28: 48–57

Deng K, Yin H, Xiong F, Feng L, Dong P, Ren M (2021) Genome-wide miRNA expression profiling in potato (Solanum tuberosum L.) reveals TOR-dependent post-transcriptional gene regulatory networks in diverse metabolic pathway. PeerJ 9: e10704

Edgar RC (2004) MUSCLE: Multiple sequence alignment with high accur-acy and high throughput. Nucleic Acids Res 32: 1792–1797

Emms DM, Kelly S (2015). OrthoFinder: solving fundamental biases in whole genome comparisons dramatically improves orthogroup inference accuracy. Genome biology 16: 157.

Esposito S, Aversano R, D’Amelia V, Villano C, Alioto D, Mirouze M, Carputo D (2018) Dicer-like and RNA-dependent RNA polymerase gene family identification and annotation in the cultivated Solanum tuberosum and its wild relative S. commersonii. Planta 248: 729–743

Fan P, Leong BJ, Last RL (2019) Tip of the trichome: evolution of acylsugar metabolic diversity in Solanaceae. Curr Opin Plant Biol 49: 8–16

Fei Q, Xia R, Meyers BC (2013) Phased, Secondary, Small Interfering RNAs in Posttranscriptional Regulatory Networks. Plant Cell 25: 2400–2415

Galindo-González L, Mhiri C, Deyholos MK, Grandbastien M-A (2017) LTR- retrotransposons in plants: Engines of evolution. Gene 626: 14–25

Goodin MM, Zaitlin D, Naidu RA, Lommel SA (2015) Nicotiana benthamiana: Its History and Future as a Model for Plant–Pathogen Interactions. Molecular Plant-Microbe Interactions 2015: 28–39

Gu M, Liu W, Meng Q, Zhang W, Chen A, Sun S, Xu G (2014) Identification of microRNAs in six solanaceous plants and their potential link with phosphate and mycorrhizal signaling. J Integr Plant Biol 56: 1164–1178

Guo Q, Qu X, Jin W (2015) PhaseTank: genome-wide computational identification of phasiRNAs and their regulatory cascades. Bioinformatics 31: 284–286

Hardigan MA, Crisovan E, Hamilton JP, Kim J, Laimbeer P, Leisner CP, Manrique-Carpintero NC, Newton L, Pham GM, Vaillancourt B, et al (2016) Genome Reduction Uncovers a Large Dispensable Genome and Adaptive Role for Copy Number Variation in Asexually Propagated Solanum tuberosum. Plant Cell 28: 388–405

Hellsten U, Wright KM, Jenkins J, Shu S, Yuan Y, Wessler SR, Schmutz J, Willis JH, Rokhsar DS (2013) Fine-scale variation in meiotic recombination in Mimulus inferred from population shotgun sequencing. Proc Natl Acad Sci U S A 110: 19478–19482

Kakrana A, Li P, Patel P, Hammond R, Anand D, Mathioni SM, Meyers BC (2017) PHASIS: A computational suite for de novo discovery and characterization of phased, siRNA-generating loci and their miRNA triggers. bioRxiv 158832

Kalyaanamoorthy S, Minh BQ, Wong TK, von Haeseler A, Jermiin LS (2017). ModelFinder: fast model selection for accurate phylogenetic estimates. Nature methods. 14:587.

Klee HJ, Giovannoni JJ (2011) Genetics and control of tomato fruit ripening and quality attributes. Annu Rev Genet 45: 41–59

Koenig DP, Sinha NR (2007) Genetic Control of Leaf Shape. Encyclopedia of Life Sciences. doi: 10.1002/9780470015902.a0020101

Kozomara A, Birgaoanu M, Griffiths-Jones S (2019) miRBase: from microRNA sequences to function. Nucleic Acids Res 47: D155–D162

Kozomara A, Griffiths-Jones S (2014) miRBase: annotating high confidence microRNAs using deep sequencing data. Nucleic Acids Res 42: D68–73

Kramerov DA, Vassetzky NS (2011) Origin and evolution of SINEs in eukaryotic genomes. Heredity 107: 487–495

Kuang H, Padmanabhan C, Li F, Kamei A, Bhaskar PB, Ouyang S, Jiang J, Buell CR, Baker B (2009) Identification of miniature inverted-repeat transposable elements (MITEs) and biogenesis of their siRNAs in the Solanaceae: new functional implications for MITEs. Genome Res 19: 42–56

Langmead B (2010) Aligning short sequencing reads with Bowtie. Curr Protoc Bioinformatics Chapter 11: Unit 11.7

Lei J, Sun Y (2014) miR-PREFeR: an accurate, fast and easy-to-use plant miRNA prediction tool using small RNA-Seq data. Bioinformatics 30: 2837–2839

Leong BJ, Lybrand DB, Lou Y-R, Fan P, Schilmiller AL, Last RL (2019) Evolution of metabolic novelty: A trichome-expressed invertase creates specialized metabolic diversity in wild tomato. Sci Adv 5: eaaw3754

Letunic I, Bork P (2019) Interactive Tree Of Life (iTOL) v4: Recent updates and new developments. Nucleic Acids Res 47: W256–W259

Liao Z, Hodén KP, Singh RK, Dixelius C (2020) Genome-wide identification of Argonautes in Solanaceae with emphasis on potato. Scientific Reports. doi: 10.1038/s41598-020-77593-y

Li H, Deng Y, Wu T, Subramanian S, Yu O (2010) Misexpression of miR482, miR1512, and miR1515 increases soybean nodulation. Plant Physiol 153: 1759– 1770

Liu Y, Teng C, Xia R, Meyers BC (2020) Phased secondary small interfering RNAs (phasiRNAs) in plants: their biogenesis, genic sources, and roles in stress responses, development, and reproduction. Plant Cell

Lunardon A, Johnson NR, Hagerott E, Phifer T (2020) Integrated annotations and analyses of small RNA–producing loci from 47 diverse plants. Genome

Martinez CC, Li S, Woodhouse MR, Sugimoto K, Sinha NR (2020) Spatial transcriptional signatures define margin morphogenesis along the proximal-distal and medio-lateral axes in tomato (Solanum lycopersicum) leaves. Plant Cell. doi: 10.1093/plcell/koaa012

Ma Z, Zhang X (2018) Actions of plant Argonautes: predictable or unpredictable? Curr Opin Plant Biol 45: 59–67

Minh BQ, Nguyen MA, von Haeseler A (2013). Ultrafast approximation for phylogenetic bootstrap. Molecular biology and evolution. 30:1188–95.

Mohorianu I, Schwach F, Jing R, Lopez-Gomollon S, Moxon S, Szittya G, Sorefan K, Moulton V, Dalmay T (2011) Profiling of short RNAs during fleshy fruit development reveals stage-specific sRNAome expression patterns: Time course study of short RNAs during fruit development. Plant J 67: 232–246

Mosher RA, Melnyk CW (2010) siRNAs and DNA methylation: seedy epigenetics. Trends in Plant Science 15: 204–210

Mueller LA, Solow TH, Taylor N, Skwarecki B, Buels R, Binns J, Lin C, Wright MH, Ahrens R, Wang Y, et al (2005) The SOL Genomics Network: a comparative resource for Solanaceae biology and beyond. Plant Physiol 138: 1310–1317

Nakano M, McCormick K, Demirci C, Demirci F, Gurazada SGR, Ramachandruni D, Dusia A, Rothhaupt JA, Meyers BC (2020) Next-Generation Sequence Databases: RNA and Genomic Informatics Resources for Plants. Plant Physiol 182: 136–146

Nguyen LT, Schmidt HA, Von Haeseler A, Minh BQ (2015). IQ-TREE: a fast and effective stochastic algorithm for estimating maximum-likelihood phylogenies. Molecular biology and evolution. 32:268–74.

Niederhuth CE, Bewick AJ, Ji L, Alabady MS, Kim KD, Li Q, Rohr NA, Rambani A, Burke JM, Udall JA, et al (2016) Widespread natural variation of DNA methylation within angiosperms. Genome Biol 17: 194

Park M, Park J, Kim S, Kwon J-K, Park HM, Bae IH, Yang T-J, Lee Y-H, Kang B-C, Choi D (2012) Evolution of the large genome in Capsicum annuum occurred through accumulation of single-type long terminal repeat retrotransposons and their derivatives. Plant J 69: 1018–1029

Pokhrel, S., Huang, K., & Meyers, B. C. (2021). Conserved and non-conserved triggers of 24-nt reproductive phasiRNAs in eudicots. BioRxiv, https://doi.org/10.1101/2021.01.20.427321

Pokhrel, S., Huang, K., Bélanger, S., Caplan, J. L., Kramer, E. M., & Meyers, B. C. (2020). Pre-meiotic, 21-nucleotide Reproductive PhasiRNAs Emerged in Seed Plants and Diversified in Flowering Plants. BioRxiv. https://doi.org/10.1101/2020.10.16.341925

Pombo MA, Rosli HG, Fernandez-Pozo N, Bombarely A (2020) Nicotiana benthamiana, A Popular Model for Genome Evolution and Plant–Pathogen Interactions. The Tobacco Plant Genome 231–247

Potato Genome Sequencing Consortium, Xu X, Pan S, Cheng S, Zhang B, Mu D, Ni P, Zhang G, Yang S, Li R, et al (2011) Genome sequence and analysis of the tuber crop potato. Nature 475: 189–195

Qin C, Yu C, Shen Y, Fang X, Chen L, Min J, Cheng J, Zhao S, Xu M, Luo Y, et al (2014) Whole-genome sequencing of cultivated and wild peppers provides insights into Capsicum domestication and specialization. Proc Natl Acad Sci U S A 111: 5135–5140

Särkinen T, Bohs L, Olmstead RG, Knapp S (2013) A phylogenetic framework for evolutionary study of the nightshades (Solanaceae): a dated 1000-tip tree. BMC Evol Biol 13: 214

Schoft VK, Chumak N, Mosiolek M, Slusarz L, Komnenovic V, Brownfield L, Twell D, Kakutani T, Tamaru H (2009) Induction of RNA-directed DNA methylation upon decondensation of constitutive heterochromatin. EMBO Rep 10: 1015–1021

Seo E, Kim T, Park JH, Yeom S-I, Kim S, Seo M-K, Shin C, Choi D (2018) Genome-wide comparative analysis in Solanaceous species reveals evolution of microRNAs targeting defense genes in Capsicum spp. DNA Res 25: 561–575

Shinozaki Y, Nicolas P, Fernandez-Pozo N, Ma Q, Evanich DJ, Shi Y, Xu Y, Zheng Y, Snyder SI, Martin LBB, et al (2018) High-resolution spatiotemporal transcriptome mapping of tomato fruit development and ripening. Nature Communications. doi: 10.1038/s41467-017-02782-9

Shivaprasad PV, Chen H-M, Patel K, Bond DM, Bruno A C, Baulcombe DC (2012) A MicroRNA Superfamily Regulates Nucleotide Binding Site–Leucine-Rich Repeats and Other mRNAs. Plant Cell 24: 859–874

Slotkin RK, Vaughn M, Borges F, Tanurdzić M, Becker JD, Feijó JA, Martienssen RA (2009) Epigenetic reprogramming and small RNA silencing of transposable elements in pollen. Cell 136: 461–472

Song Q-X, Liu Y-F, Hu X-Y, Zhang W-K, Ma B, Chen S-Y, Zhang J-S (2011) Identification of miRNAs and their target genes in developing soybean seeds by deep sequencing. BMC Plant Biol 11: 5

Sun W, Xiang X, Zhai L, Zhang D, Cao Z, Liu L, Zhang Z (2018) AGO18b negatively regulates determinacy of spikelet meristems on the tassel central spike in maize. J Integr Plant Biol 60: 65–78

Sun X, Zhang J, Li A, Yuan X (2013) mirPD: A pattern-based approach for identifying microRNAs from deep sequencing data. Digital Signal Processing 23: 1887–1896

Taller D, Bálint J, Gyula P, Nagy T, Barta E, Baksa I, Szittya G, Taller J, Havelda Z (2018) Correction: Expansion of Capsicum annum fruit is linked to dynamic tissue-specific differential expression of miRNA and siRNA profiles. PLoS One 13: e0203582

Tomato Genome Consortium (2012) The tomato genome sequence provides insights into fleshy fruit evolution. Nature 485: 635–641

Turner SD (2014) qqman: an R package for visualizing GWAS results using Q-Q and manhattan plots. bioRxiv 005165

Vaucheret H, Vazquez F, Crété P, Bartel DP (2004) The action of ARGONAUTE1 in the miRNA pathway and its regulation by the miRNA pathway are crucial for plant development. Genes Dev 18: 1187–1197

Verhoeven KJF, Preite V (2014) Epigenetic variation in asexually reproducing organisms. Evolution 68: 644–655

Vicient CM, Casacuberta JM (2017) Impact of transposable elements on polyploid plant genomes. Ann Bot 120: 195–207

Vrbsky J, Akimcheva S, Watson JM, Turner TL, Daxinger L, Vyskot B, Aufsatz W, Riha K (2010) siRNA-mediated methylation of Arabidopsis telomeres. PLoS Genet 6: e1000986

de Vries S, Kloesges T, Rose LE (2015) Evolutionarily Dynamic, but Robust, Targeting of Resistance Genes by the miR482/2118 Gene Family in the Solanaceae. Genome Biol Evol 7: 3307–3321

Waese J, Fan J, Pasha A, Yu H, Fucile G, Shi R, Cumming M, Kelley L, Sternberg M, Krishnakumar V, Ferlanti E, Miller J, Town C, Stuerzlinger W, Provart NJ (2017). ePlant: Visualizing and Exploring Multiple Levels of Data for Hypothesis Generation in Plant Biology. Plant Cell 29:1806–1821.

Wang X, Weigel D, Smith LM (2013) Transposon variants and their effects on gene expression in Arabidopsis. PLoS Genet 9: e1003255

Wang Z, Hardcastle TJ, Pastor AC, Yip WH, Tang S, Baulcombe DC (2018) A novel DCL2-dependent miRNA pathway in tomato affects susceptibility to RNA viruses. Genes & Development 32: 1155–1160

Won SY, Yumul RE, Chen X (2014) Small RNAs in Plants. In SH Howell, ed, Molecular Biology. Springer New York, New York, NY, pp 95–127

Wu, H., Li, B., Iwakawa, H. oki, Pan, Y., Tang, X., Ling-hu, Q., Liu, Y., Sheng, S., Feng, L., Zhang, H., Zhang, X., Tang, Z., Xia, X., Zhai, J., & Guo, H. (2020). Plant 22-nt siRNAs mediate translational repression and stress adaptation. Nature 581: 81–93

Wu L, Zhang Q, Zhou H, Ni F, Wu X, Qi Y (2009) Rice MicroRNA Effector Complexes and Targets. The Plant Cell 21: 3421–3435

Xia R, Meyers BC, Liu Z, Beers EP, Ye S, Liu Z (2013) MicroRNA superfamilies descended from miR390 and their roles in secondary small interfering RNA Biogenesis in Eudicots. Plant Cell 25: 1555–1572

Xia R, Xu J, Arikit S, Meyers BC (2015) Extensive Families of miRNAs and PHAS Loci in Norway Spruce Demonstrate the Origins of Complex phasiRNA Networks in Seed Plants. Mol Biol Evol 32: 2905–2918

Xie Z, Kasschau KD, Carrington JC (2003) Negative feedback regulation of Dicer-Like1 in Arabidopsis by microRNA-guided mRNA degradation. Curr Biol 13: 784– 789

Zhai J, Jeong D-H, De Paoli E, Park S, Rosen BD, Li Y, González AJ, Yan Z, Kitto SL, Grusak MA, et al (2011) MicroRNAs as master regulators of the plant NB-LRR defense gene family via the production of phased, trans-acting siRNAs. Genes Dev 25: 2540–2553

Zhang H, Xia R, Meyers BC, Walbot V (2015) Evolution, functions, and mysteries of plant ARGONAUTE proteins. Curr Opin Plant Biol 27: 84–90

Zhang R, Marshall D, Bryan GJ, Hornyik C (2013) Identification and characterization of miRNA transcriptome in potato by high-throughput sequencing. PLoS One 8: e57233

Zuo J, Grierson D, Courtney LT, Wang Y, Gao L, Zhao X, Zhu B, Luo Y, Wang Q, Giovannoni JJ (2020) Relationships between genome methylation, levels of non-coding RNAs, mRNAs and metabolites in ripening tomato fruit. Plant J 103: 980–994

