## Supplemental Figures 1-7 for "The Evolutionary History of Small RNAs in the Solanaceae"

**Supplemental Figure 1: Phylogenetic analysis of proteins involved in sRNA biogenesis.** Phylogenetic tree showing gene family relationships for *DRB* (A), *NPR* (B), *RDR* (C)  in the Solanaceae. Subclades of the protein families are drawn with distinct colors. Orthologous groups were identified using OrthoFinder and SonicParanoid. Protein alignments and phylogenetic analyses were performed using MUSCLE and IQ- TREE. Phylogenetic tree was visualized using iTOL.

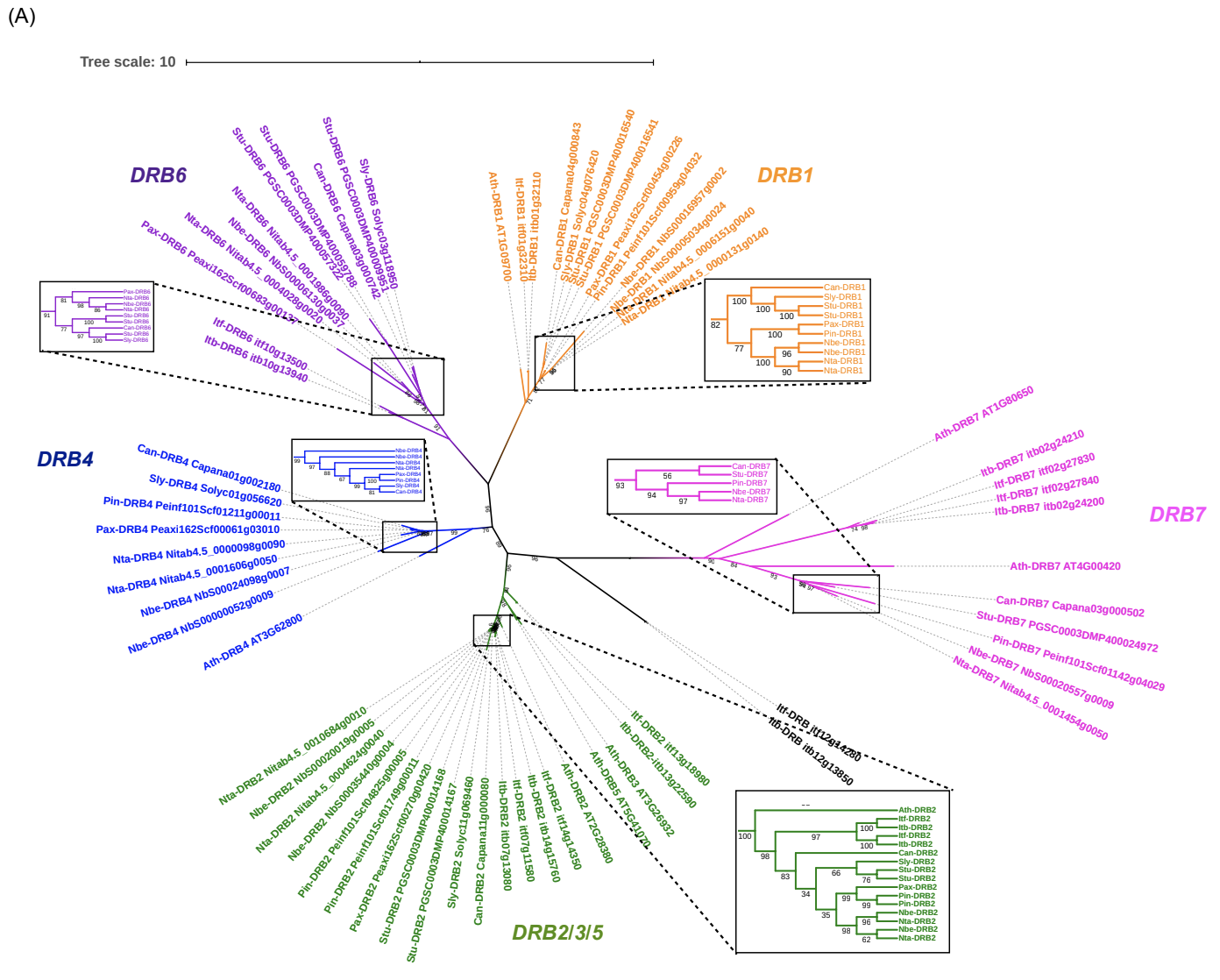

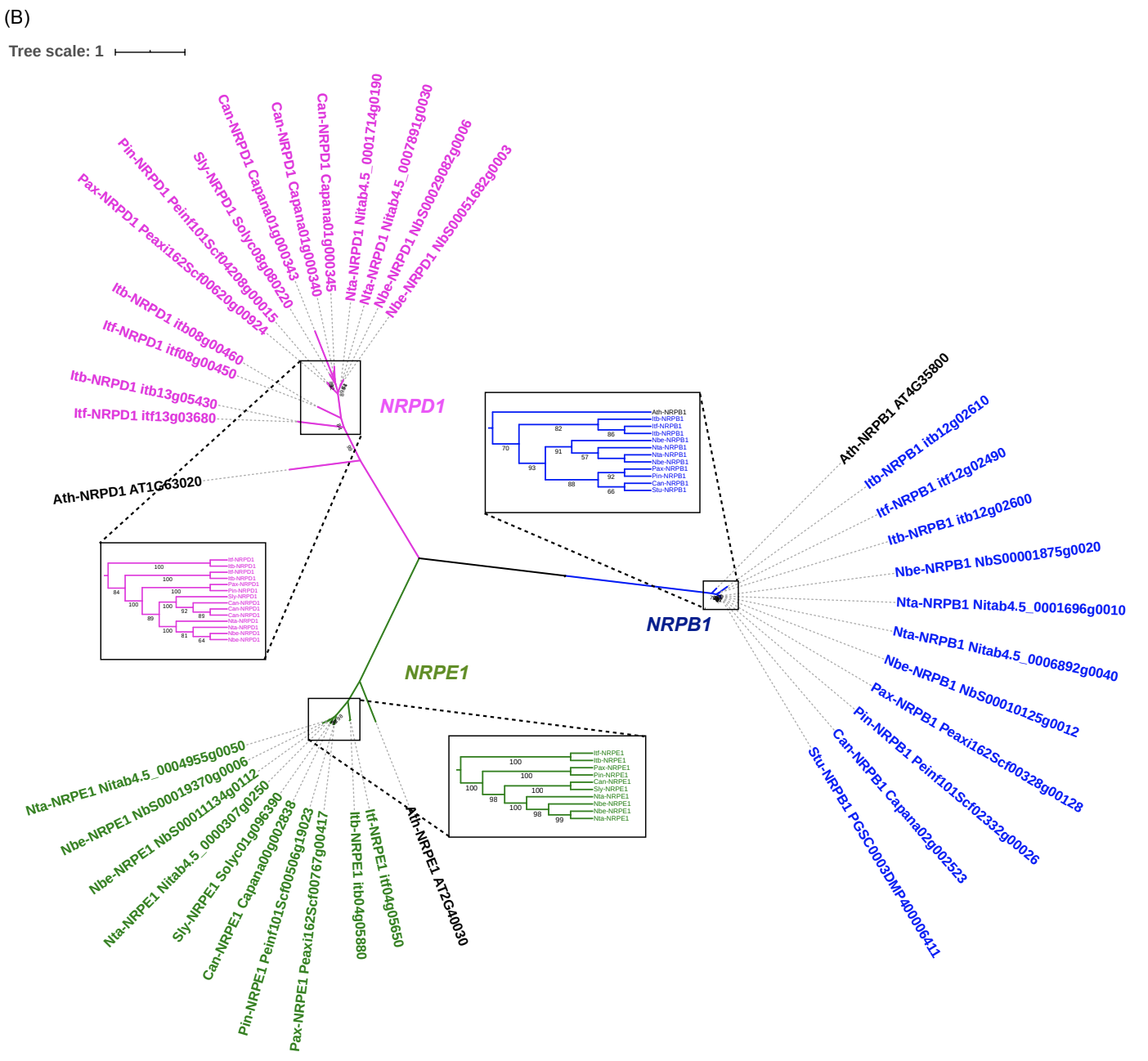

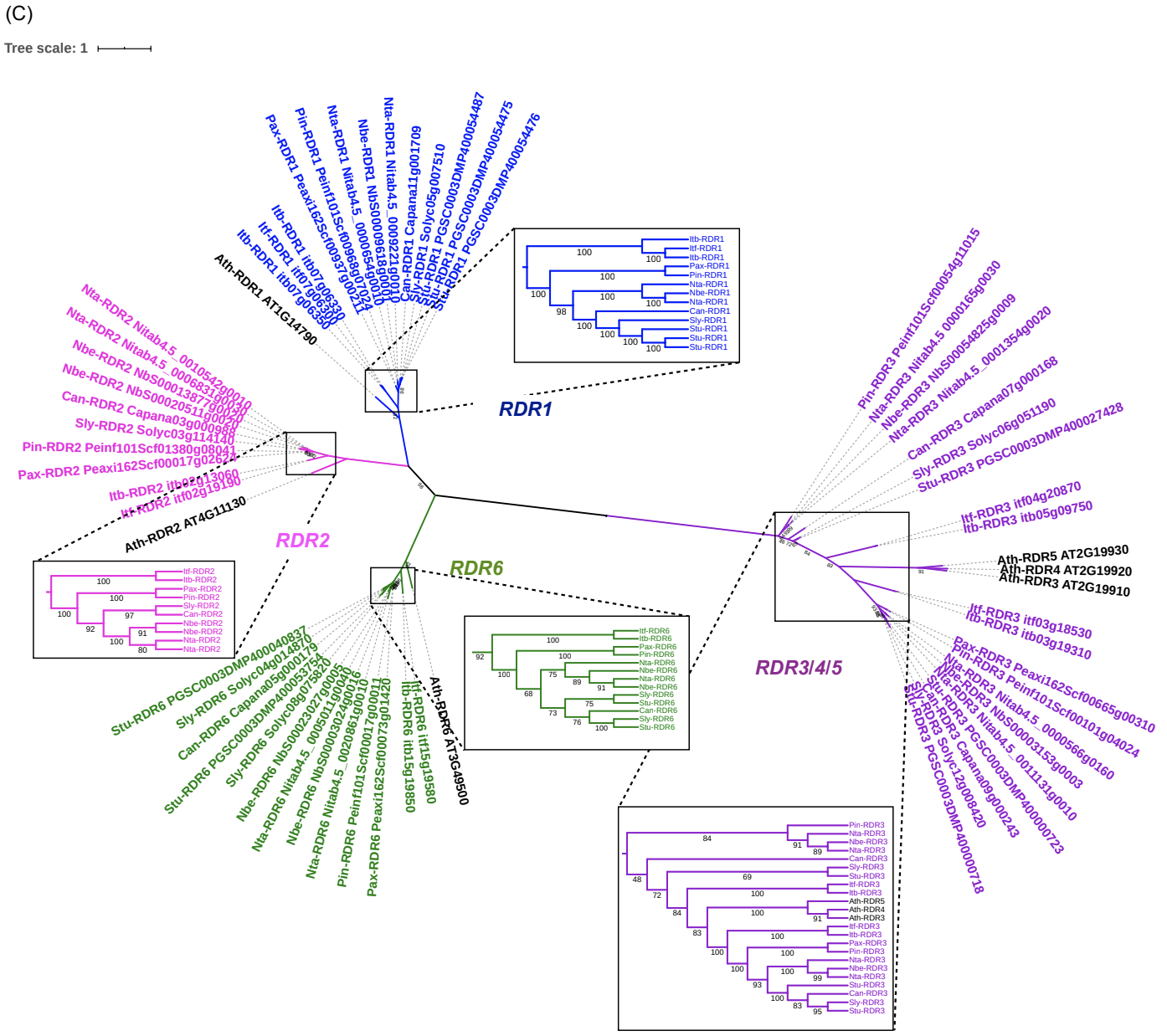

**Supplemental Figure 2: Expression of novel versus conserved miRNAs across the Solanaceae.** Novel miRNAs are expressed at significantly lower levels than miRBase conserved miRNAs based on two-tailed T-tests between novel and conserved miRNAs for each species: benthi P-value = 0.00057903, pepper P-value = 1.98548E-08, potato P-value = 1.97921E-06, tomato P-value = 2.27413E-07, petunia P-value = 2.39254E-05, tobacco P-value = 5.89638E-08.

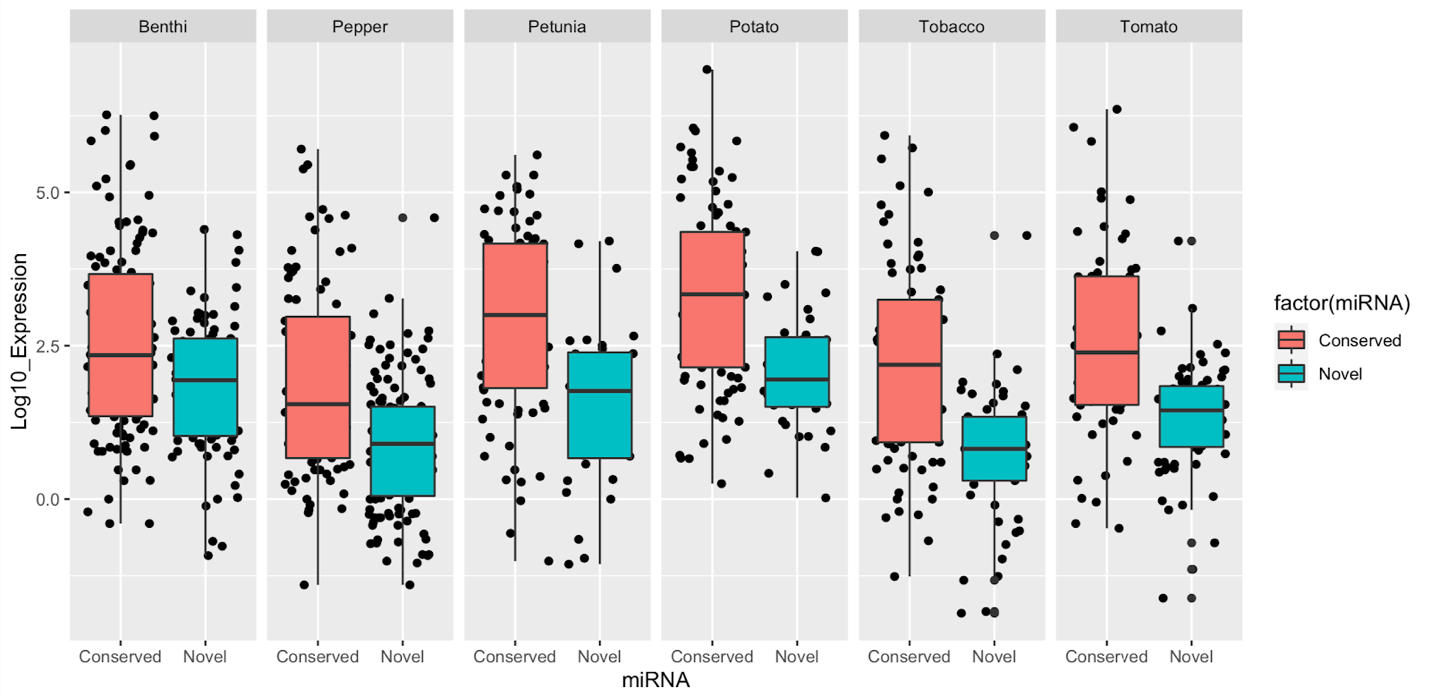

**Supplemental Figure 3: Cross-species alignment of miR5303.** Comparative sequence alignment of miR5303 across Solanaceae species reveals a highly-conserved core sequence for most variants. There are a small number of entries from Nicotiana and Petunia species with unrelated sequence identity that are likely the result of mis-annotated miRbase entries.

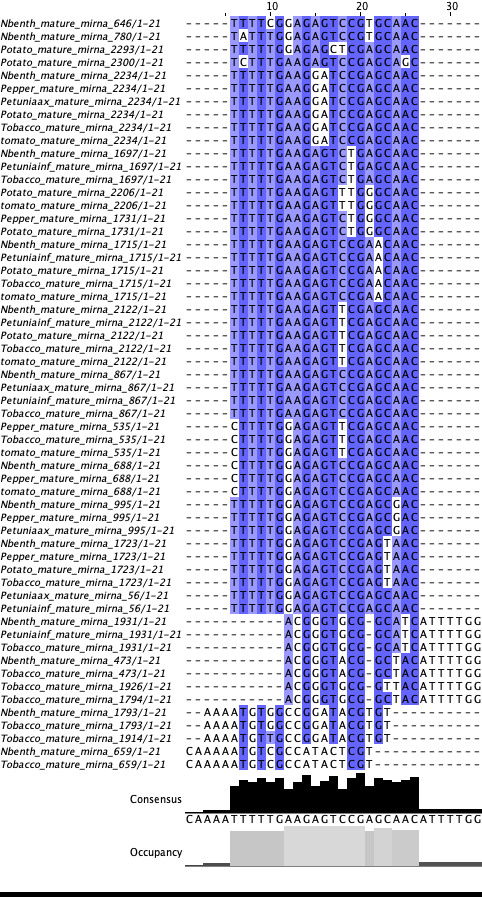

**Supplemental Figure 4: Conserved *Nicotiana* and *Solanum* TAS loci.**

Conserved *TASL1/2* loci in tobacco (A) and benthi (B) show strong phasing scores and very high sequence conservation (C). Dominant 21-nt siRNAs produced by the *TASL1/2* loci align with perfect/near perfect matches to PPR targets in the benthi genome (D). Benthi expresses two additional PHAS loci that are targeted by miR7122 (E and F), but these loci do not show high sequence conservation with the tobacco *TASL1/2* locus. A conserved *Solanum* TAS locus on Chromosome 5 that is expressed in Tomato (G) and Potato (H), shows 83% cross-species sequence conservation (I), and produces an abundant 21-nt siRNA that has sequence complementarity to MYB TFs (J).

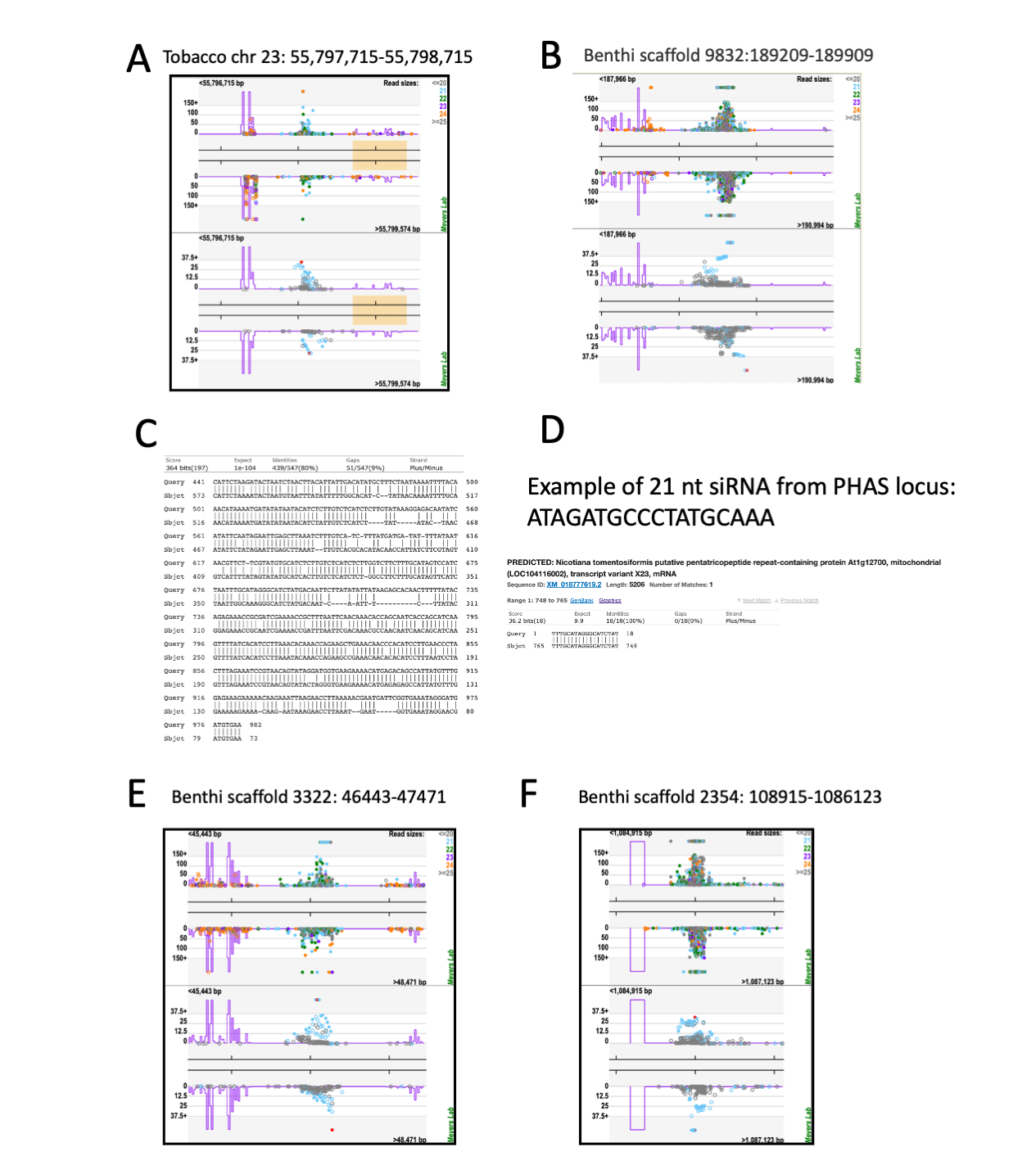

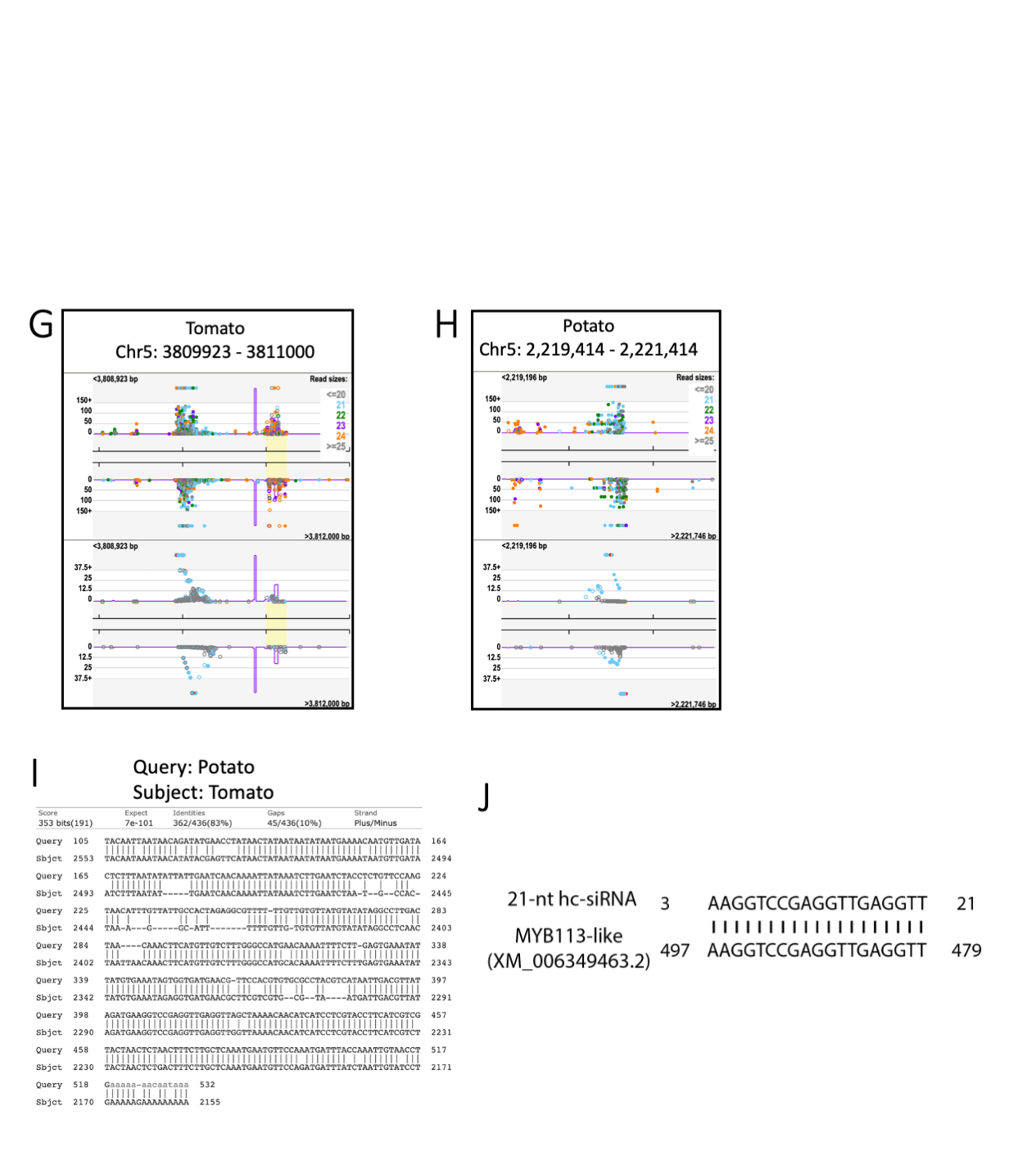

**Supplemental Figure 5: LTR elements are the most abundant Transposable Elements (TEs) in the analyzed Solanacea genomes, while SINEs are the element producing a higher amount of heterochromatic small RNAs (hc-siRNAs).** Histograms represent the percentage of each TE superfamily for the 6 study genomes (A), as well as the percentage of hc-siRNAs derived from each TE superfamily (B). The x-axis represents the different species and the y-axis represents the percentage of each category.

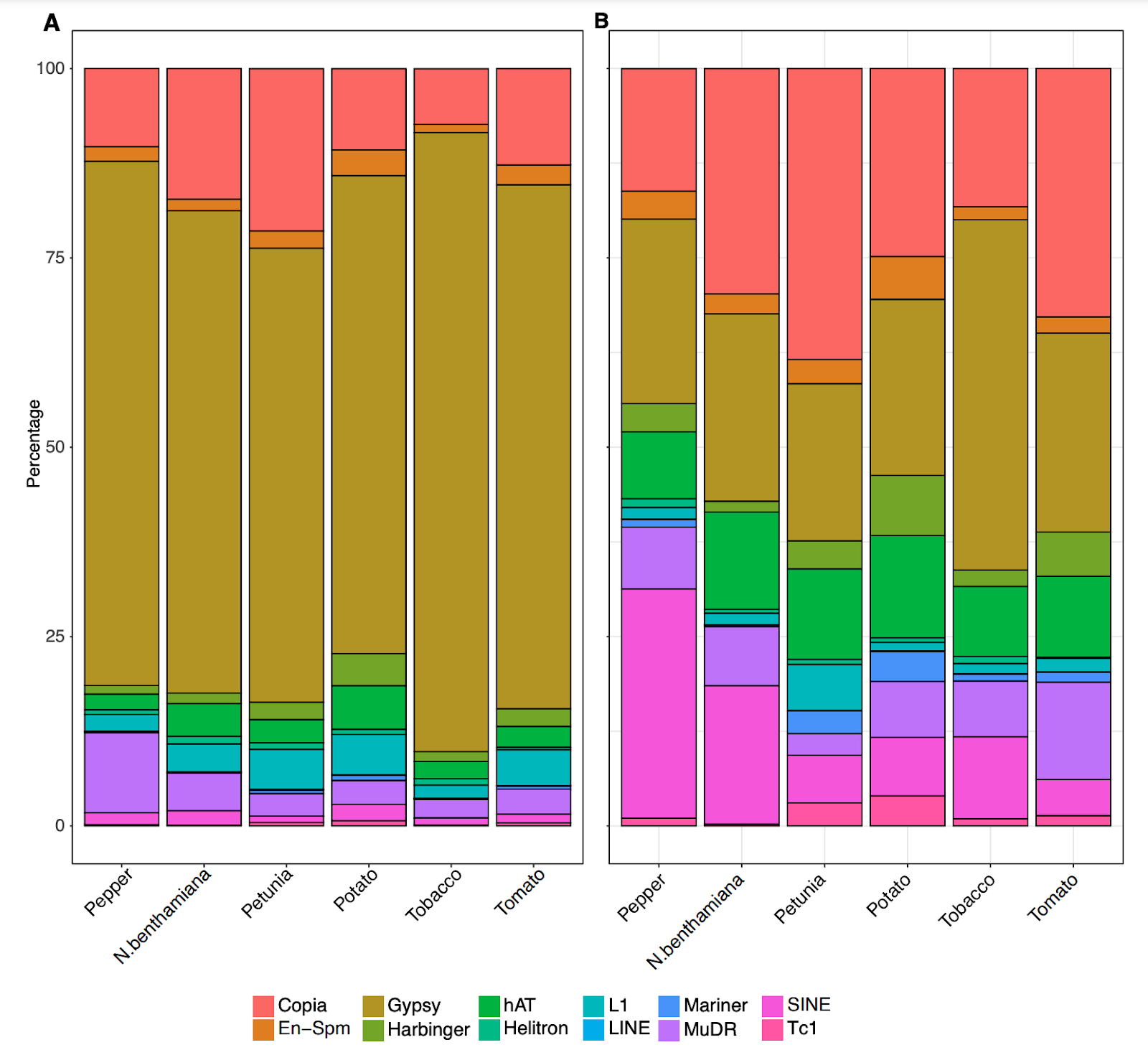

**Supplemental Figure 6: Expression of filtered 22nt sRNA clusters.**

(A) Heatmap showing the expression of the 390 clusters in the four samples. Visible virus-induced and virus-repressed clusters are marked by a red and blue line, respectively. (B) Dot plot of the logarithmic expression fold change of infected vs. control samples. Thresholds are shown in red and blue dashed lines, respectively, and the clusters passing the thresholds are highlighted as red and blue dots (no cluster), respectively. Note that infinity log2 fold change is plotted as some value above 15.

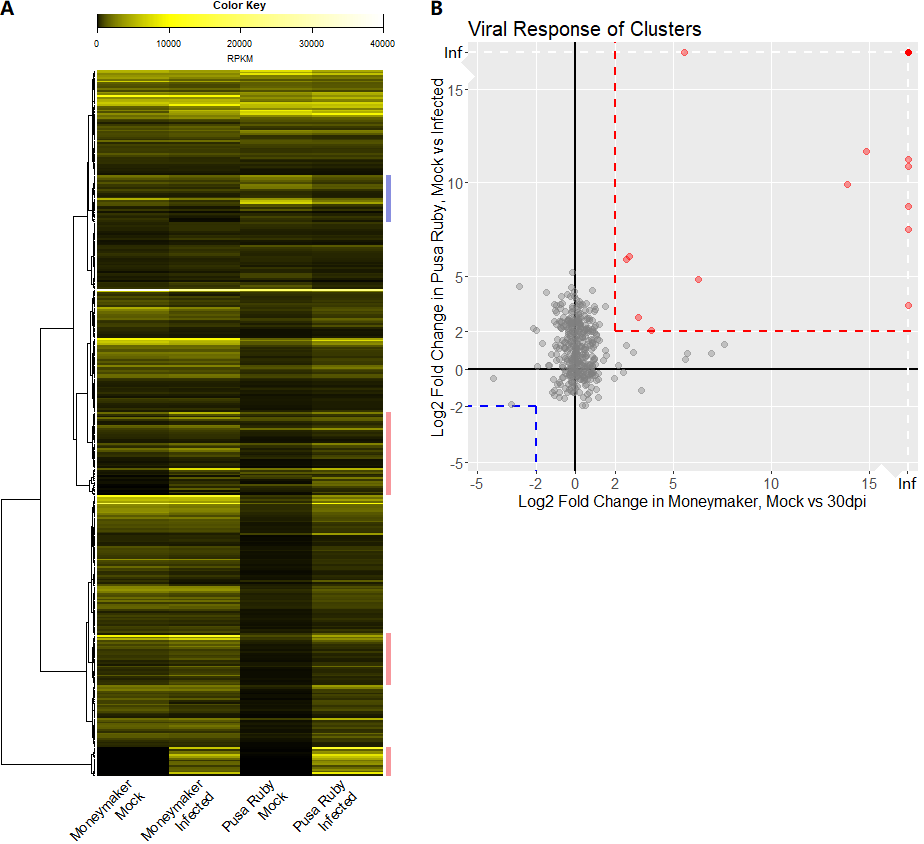

**Supplemental Figure 7: Size distribution plots for all of the sRNA libraries included in this study.** Reads were trimmed to 35 nucleotides in length and abundance in reads per million (RPM) is plotted for an 18-35 nucleotide size range. Libraries lacking enrichment in 21 and 24 nt size distributions were considered low quality and removed from this study. (A) nbenthi libraries, (B) tobacco, (C) petunia, (D) pepper, (E) tomato, and (F) potato.

A) Nicotiana

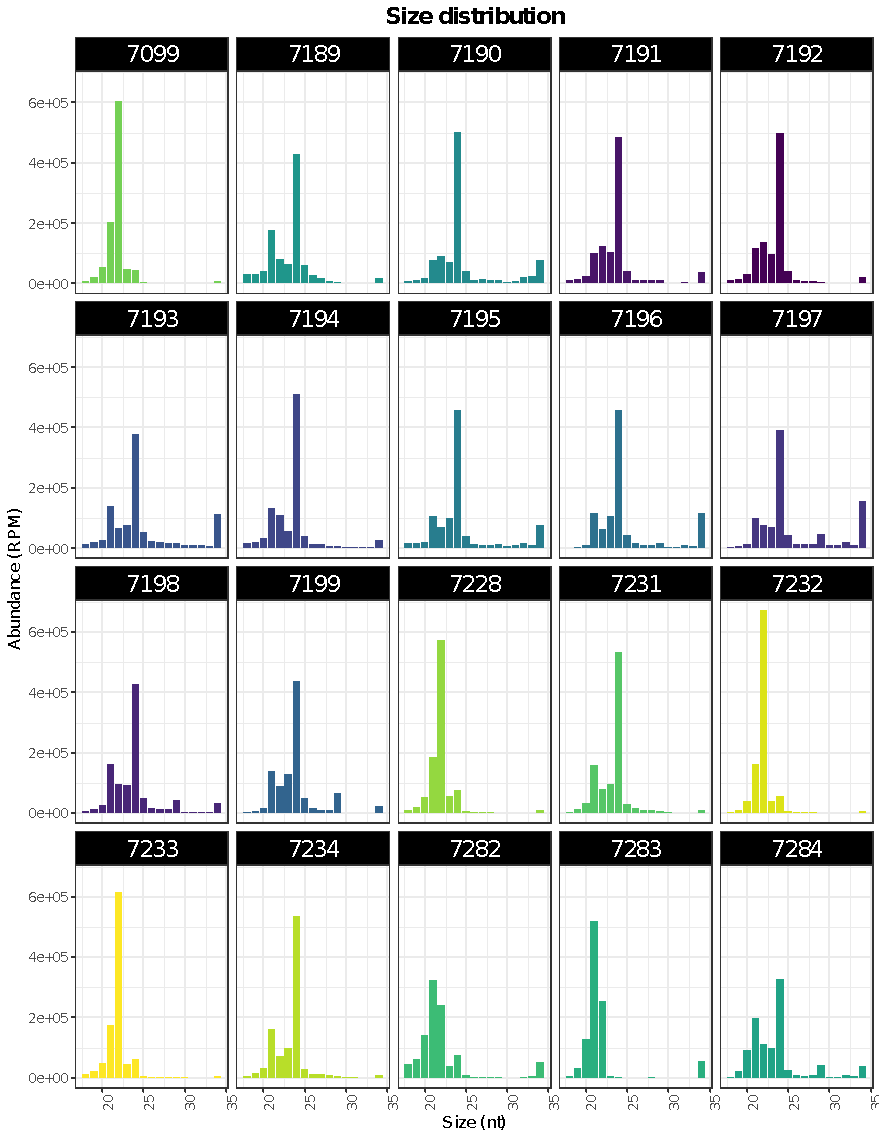

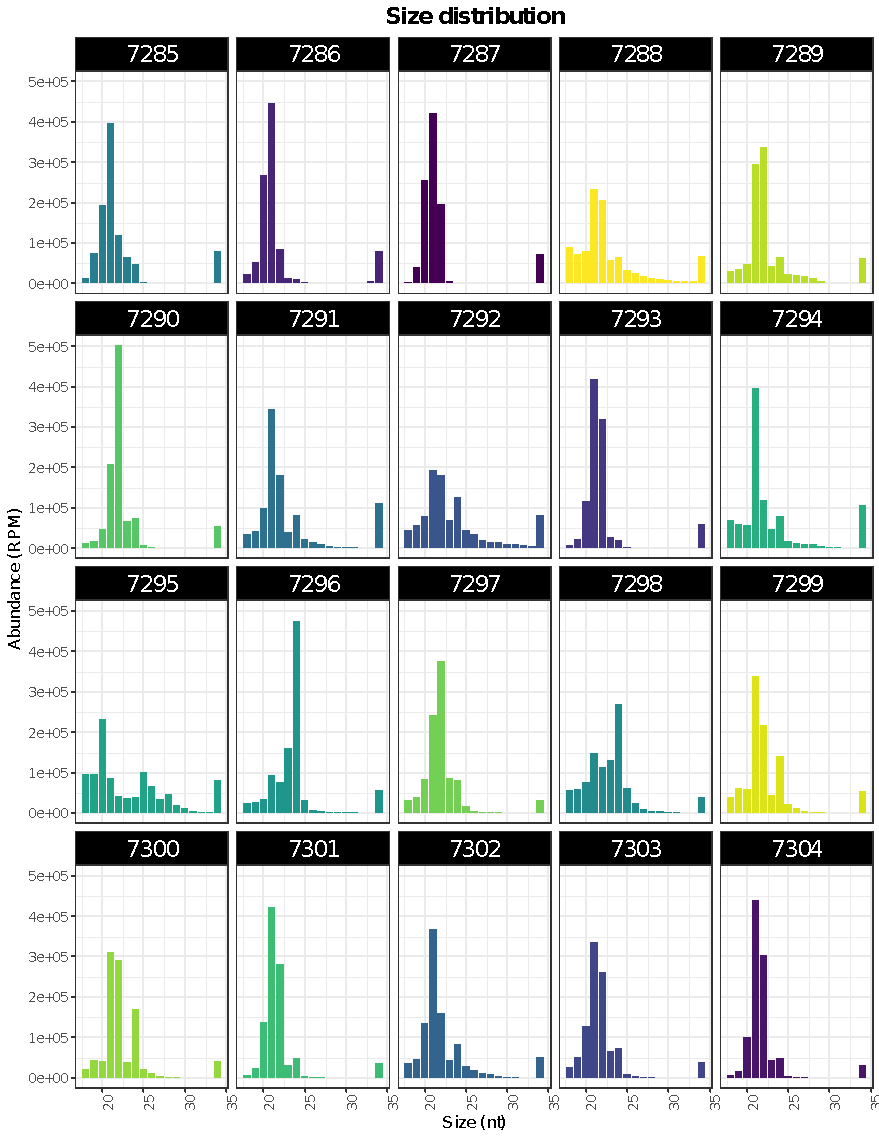

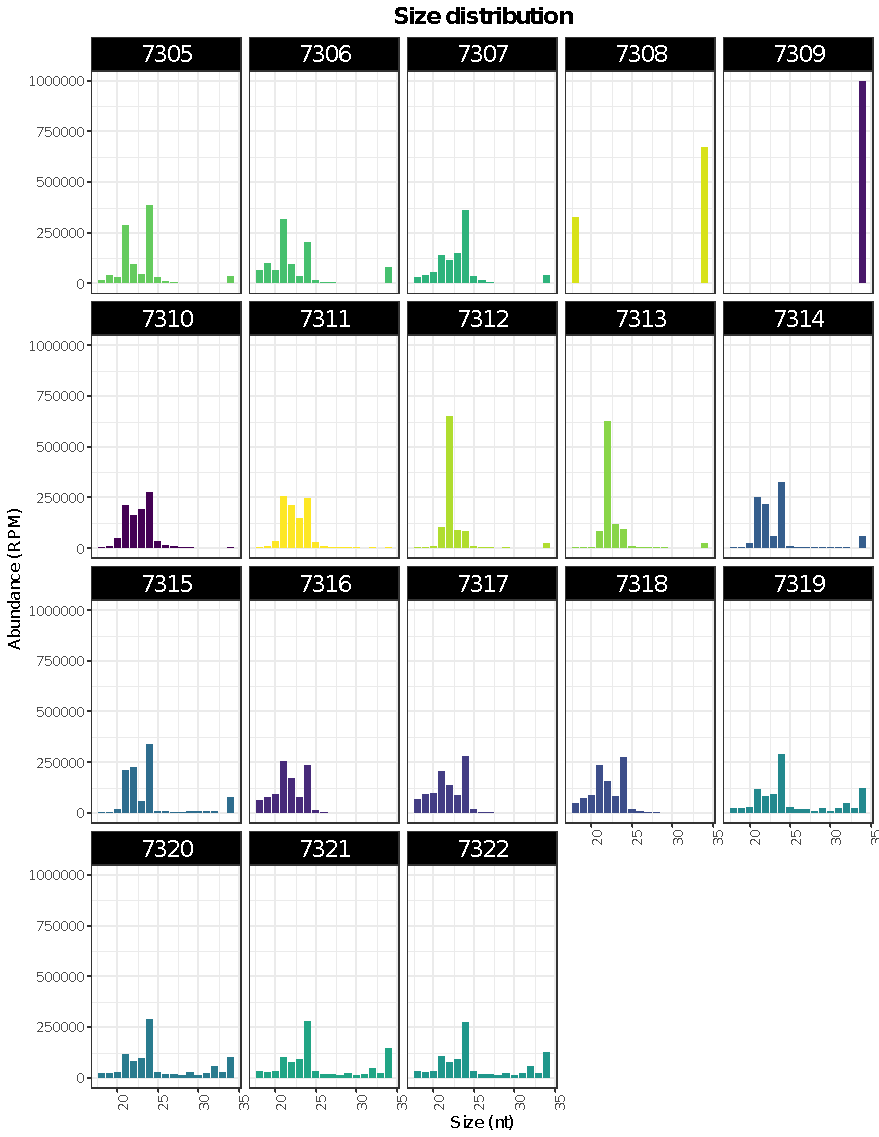

(B) tobacco

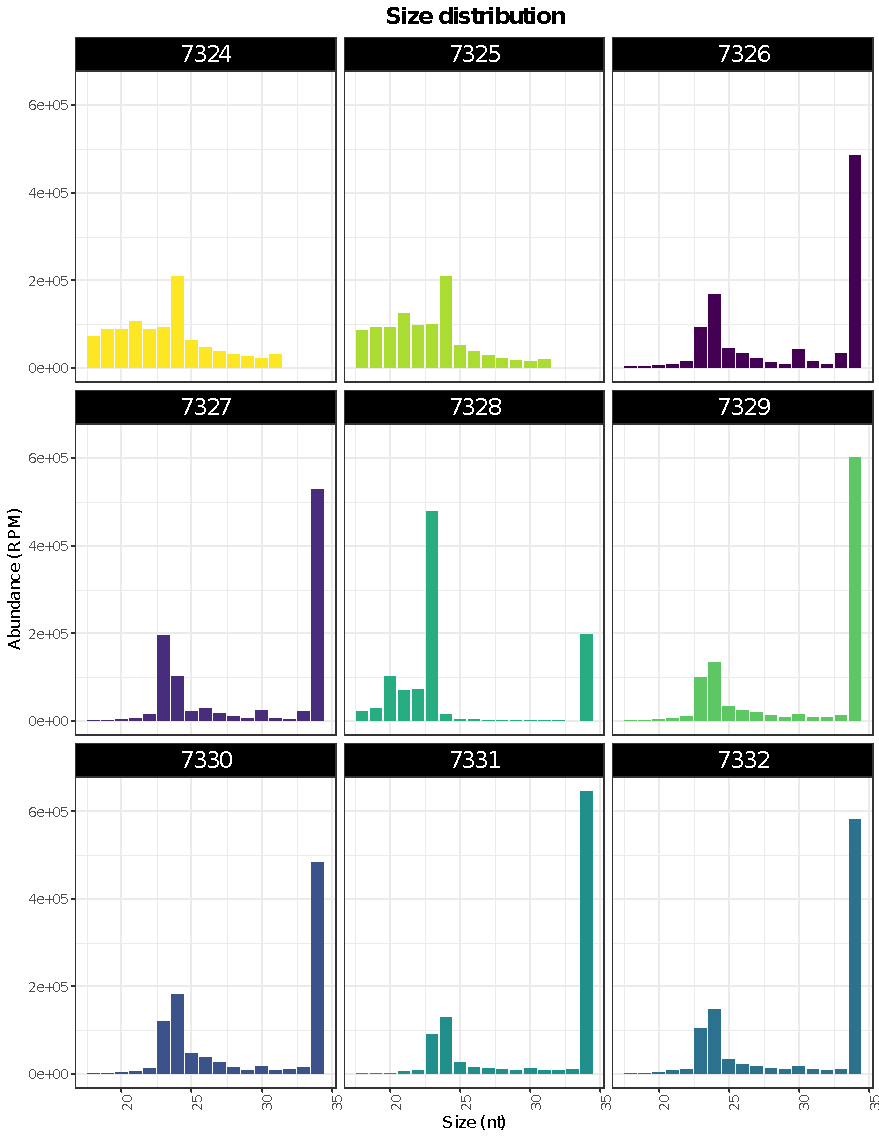

(C) petunia

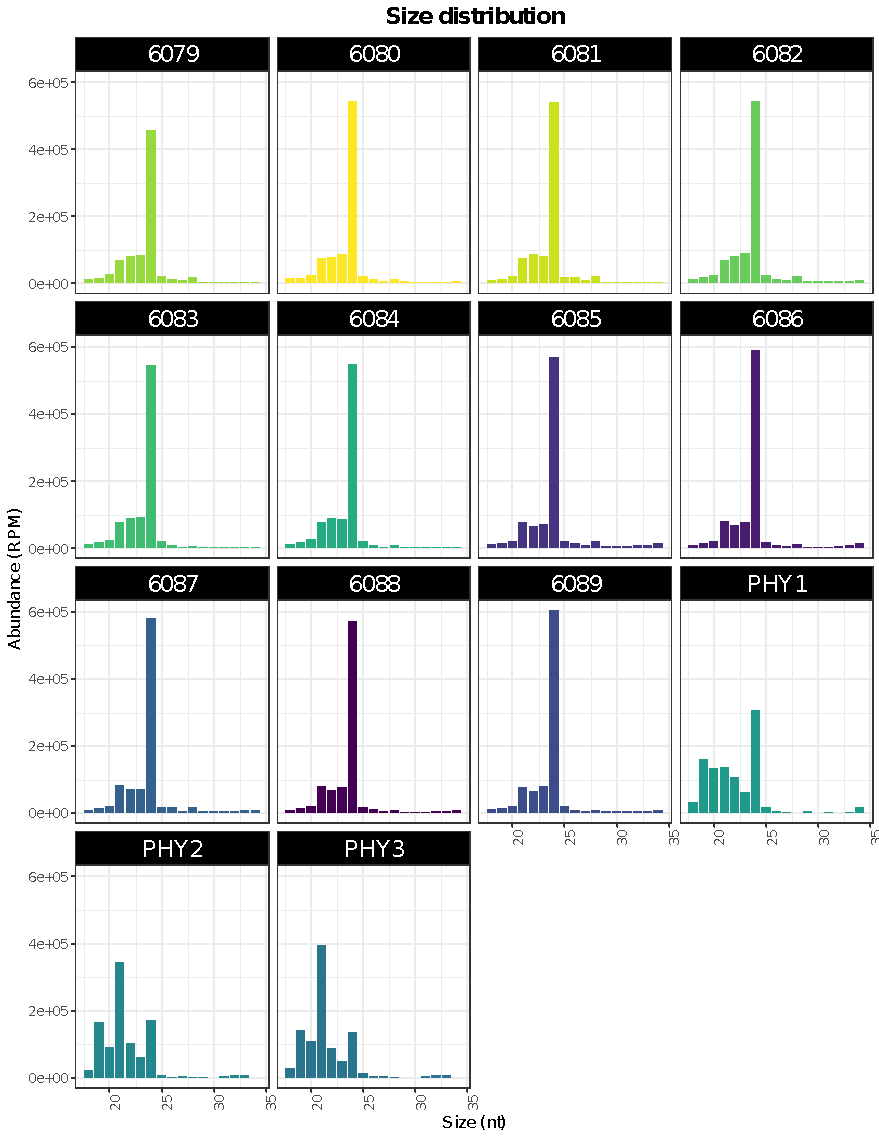

(D) pepper

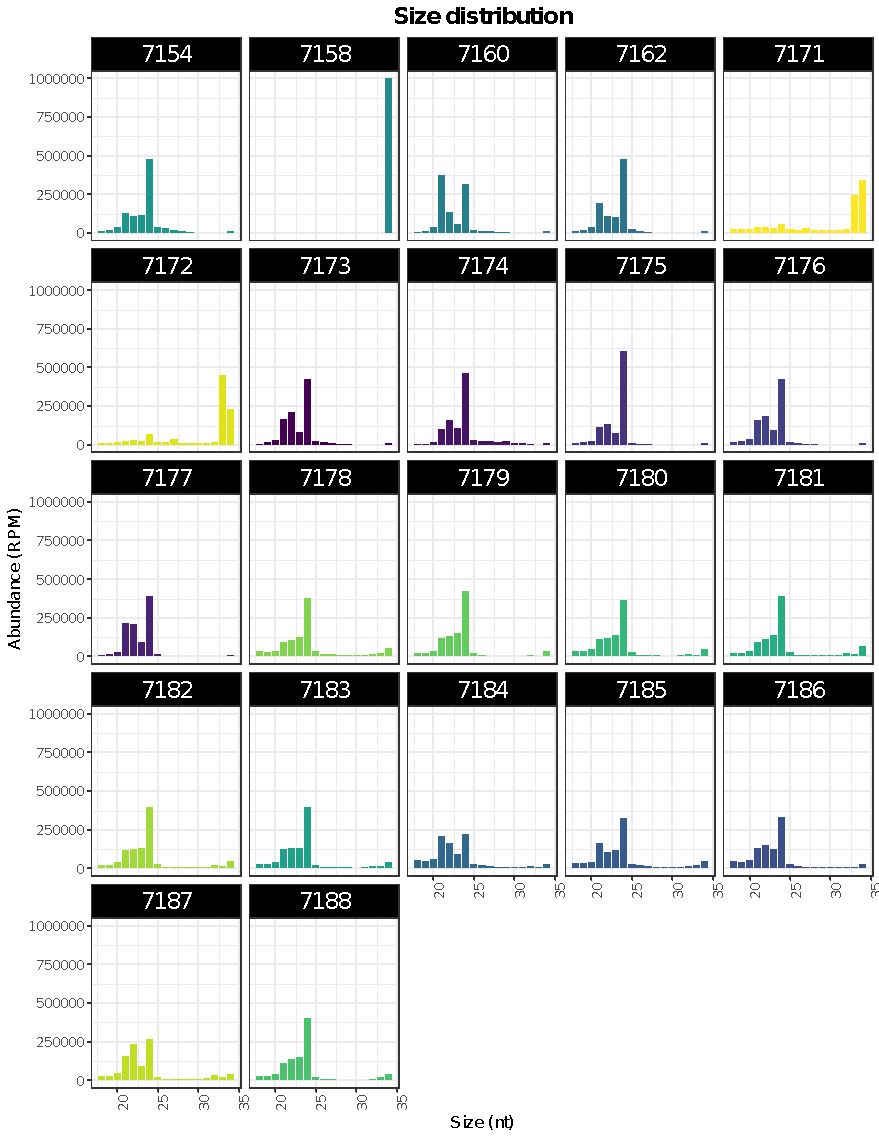

(E) tomato

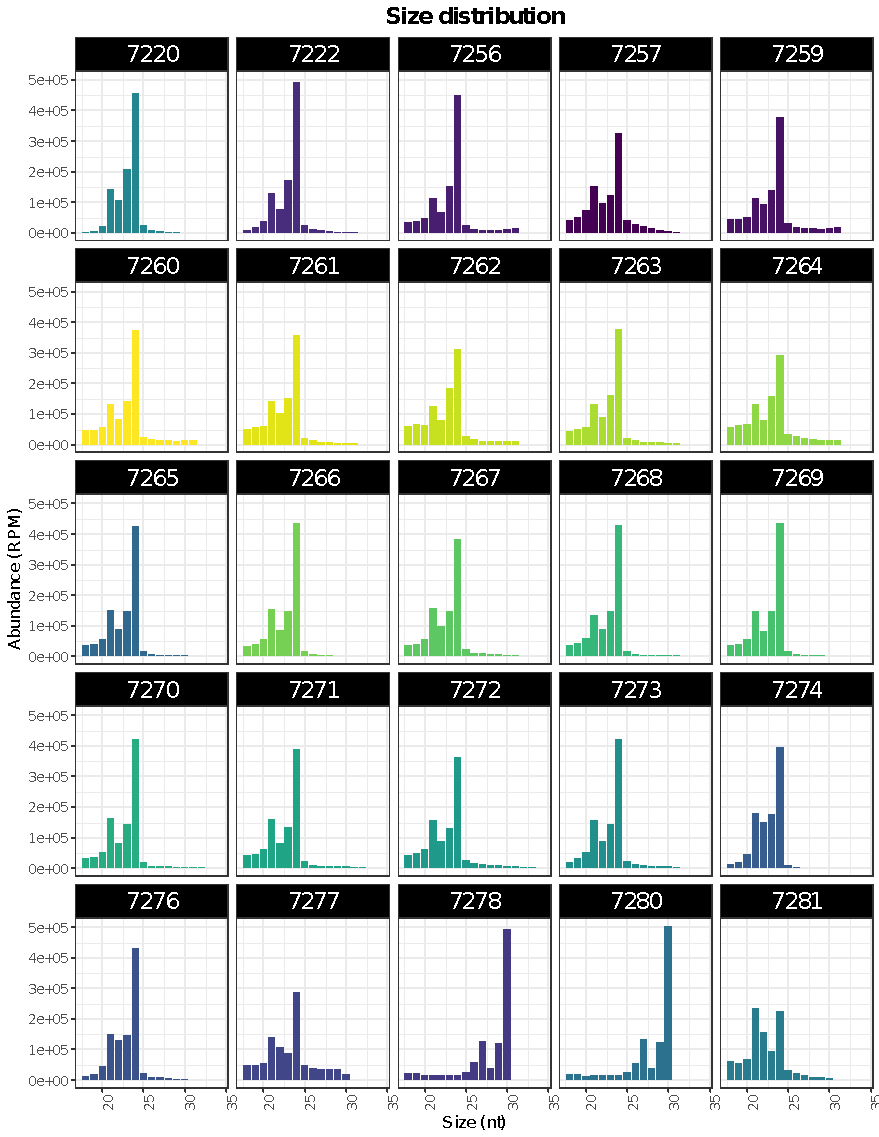

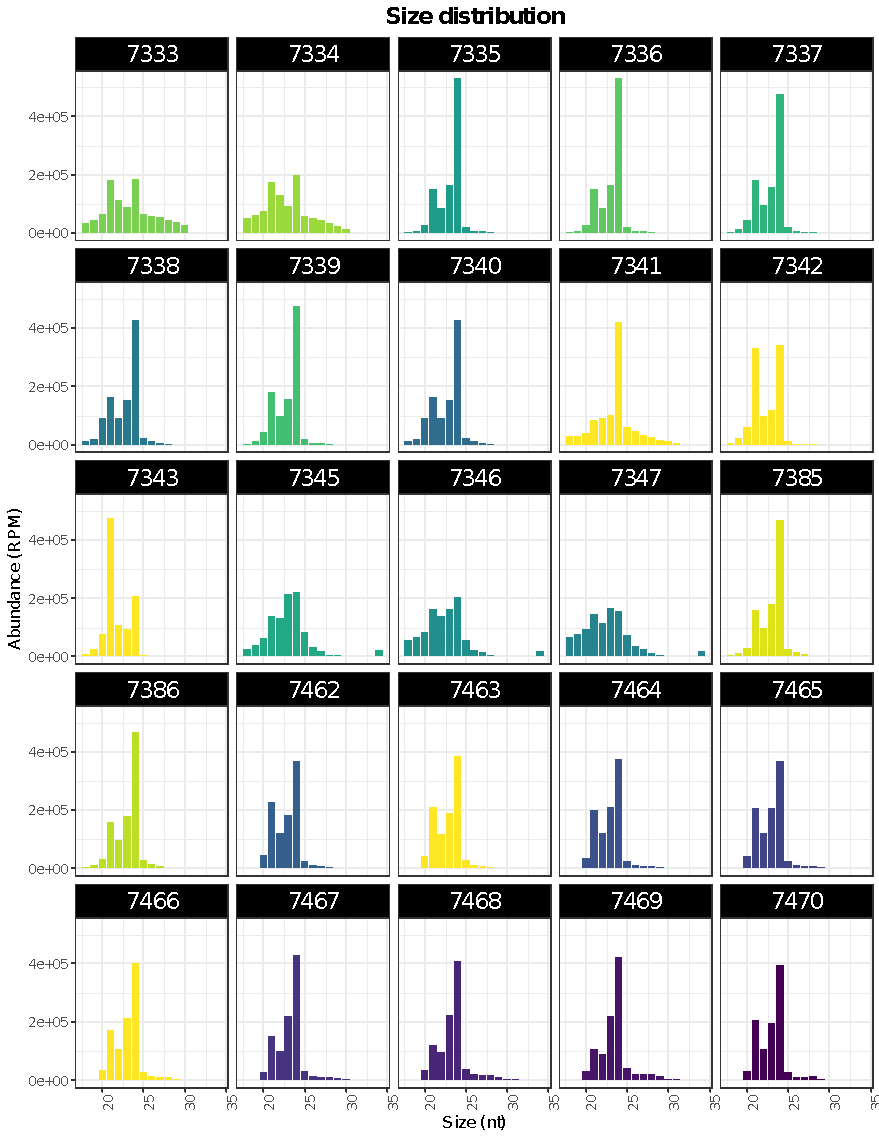

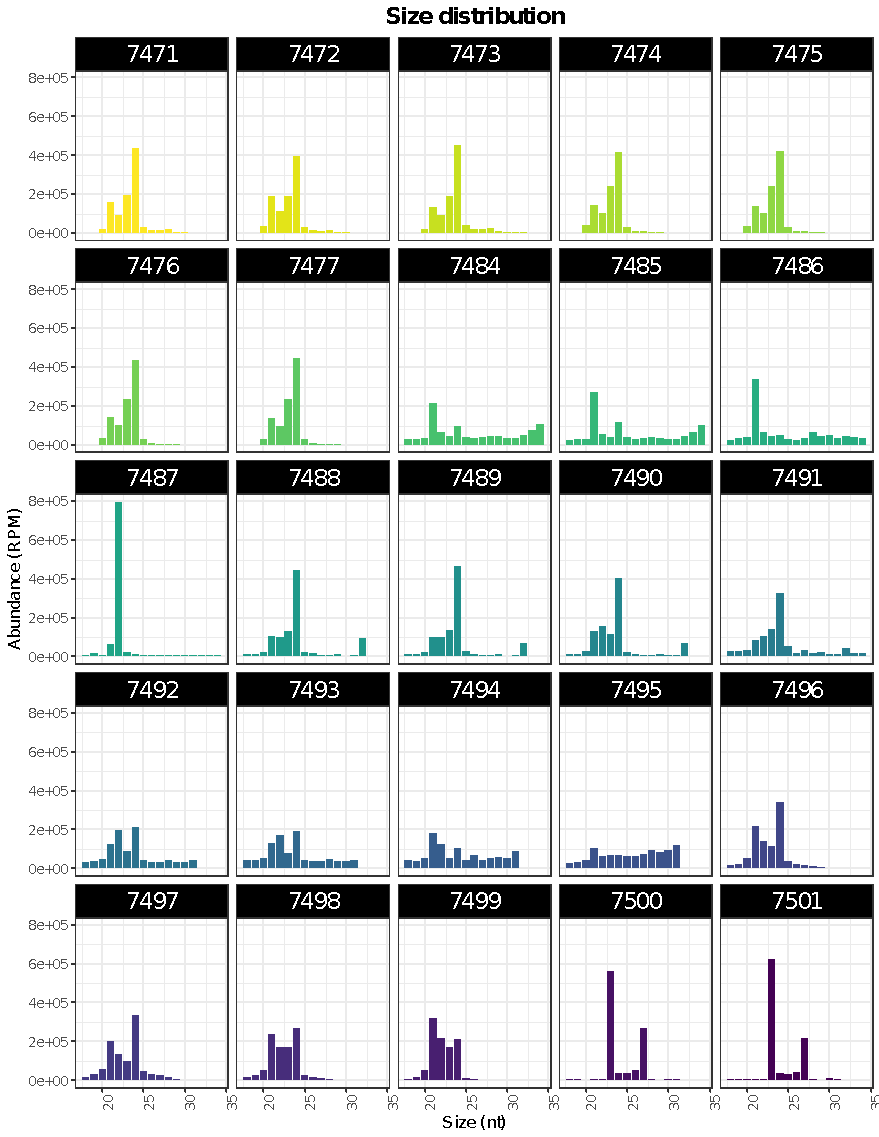

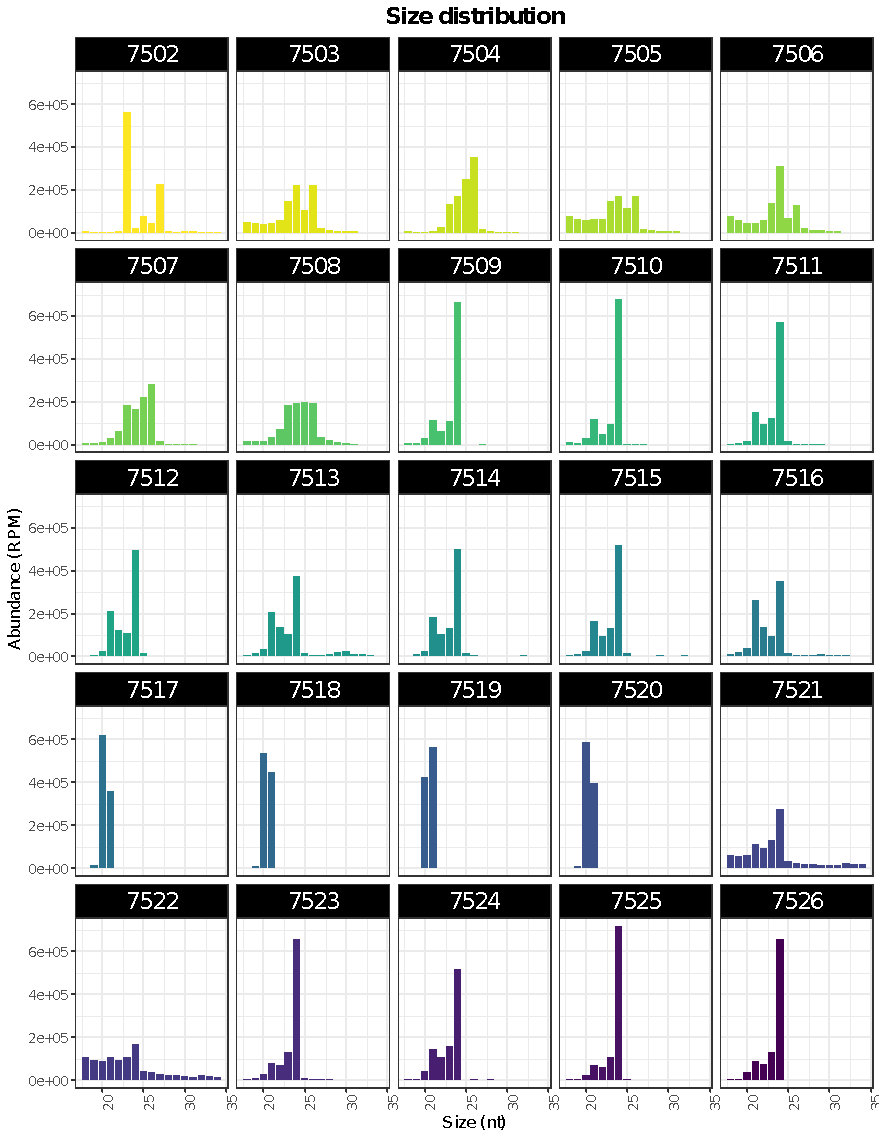

(F) potato

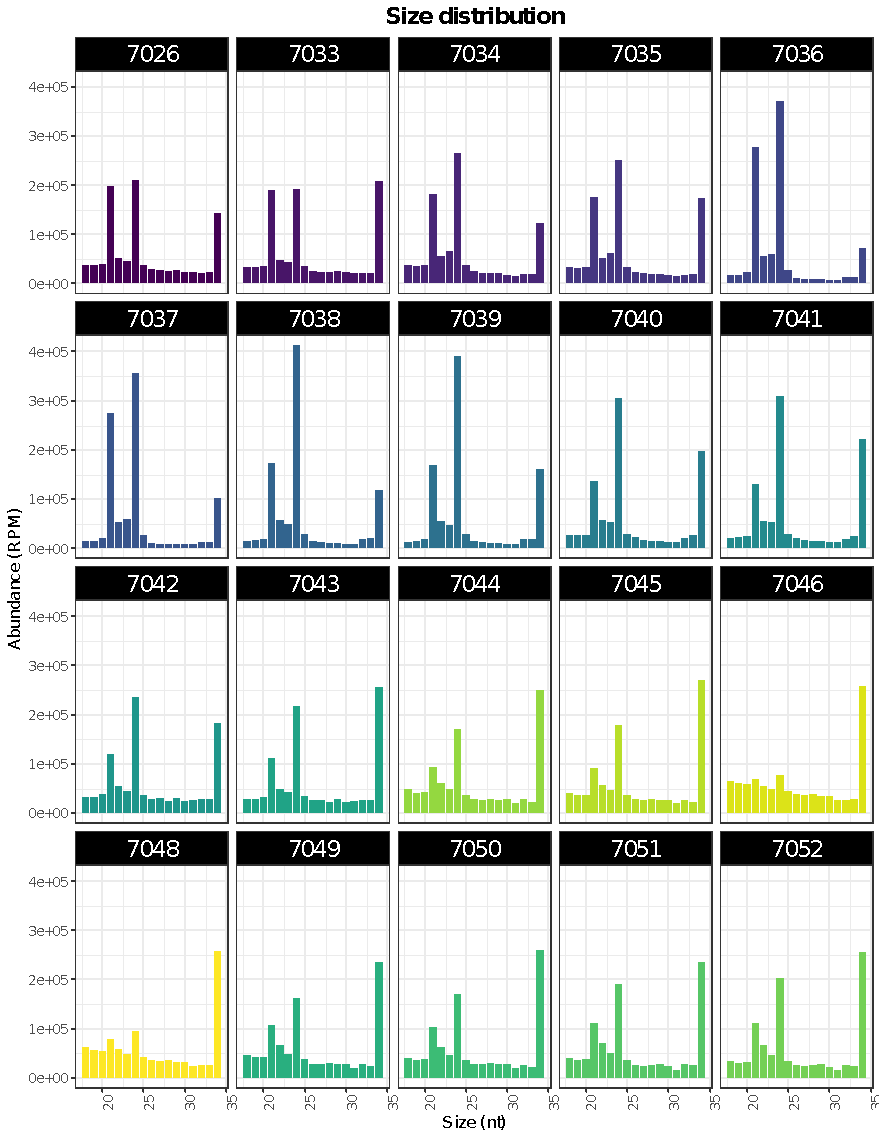

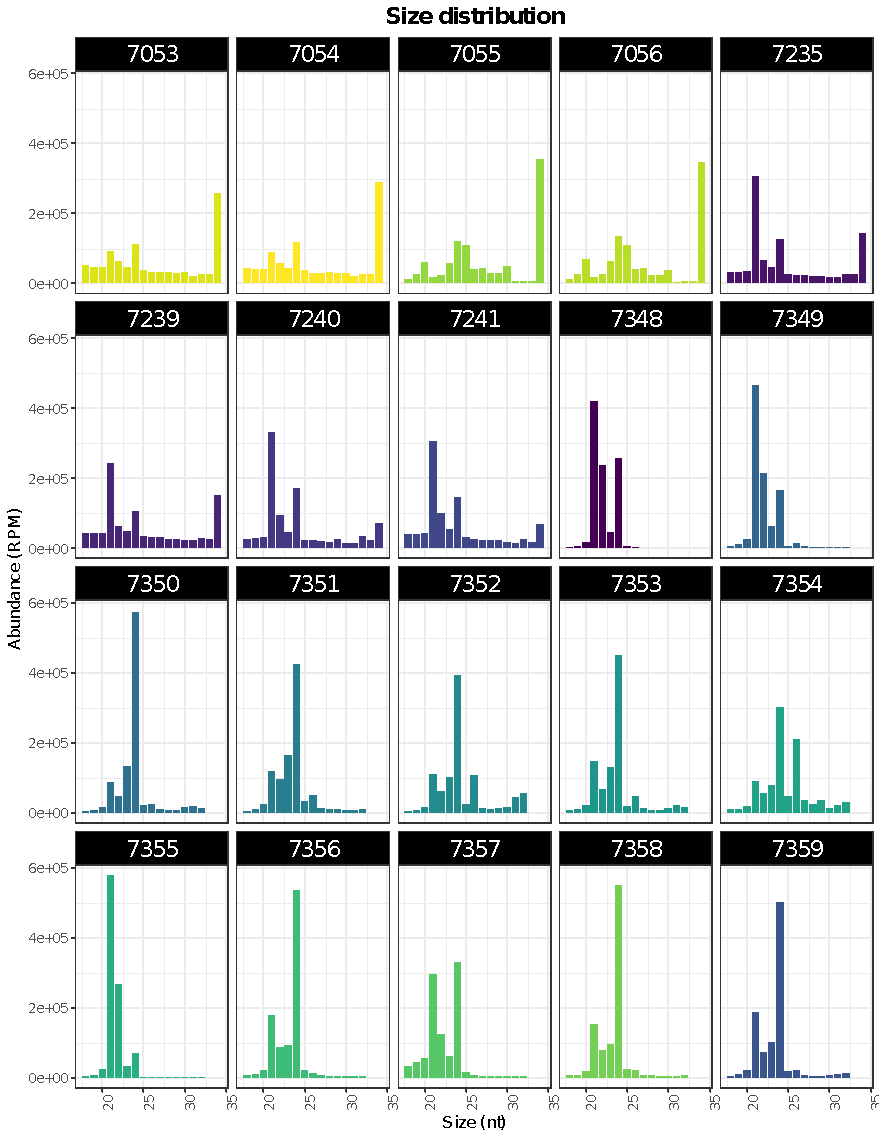

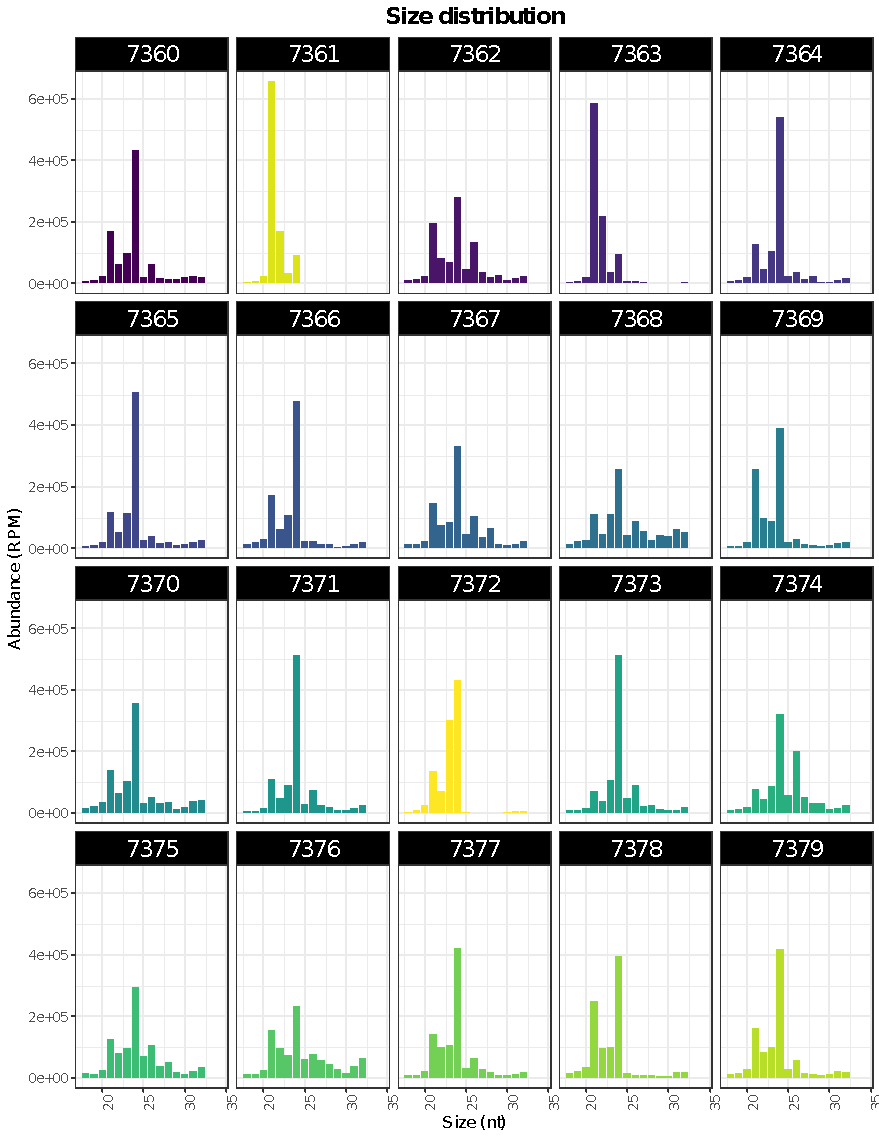

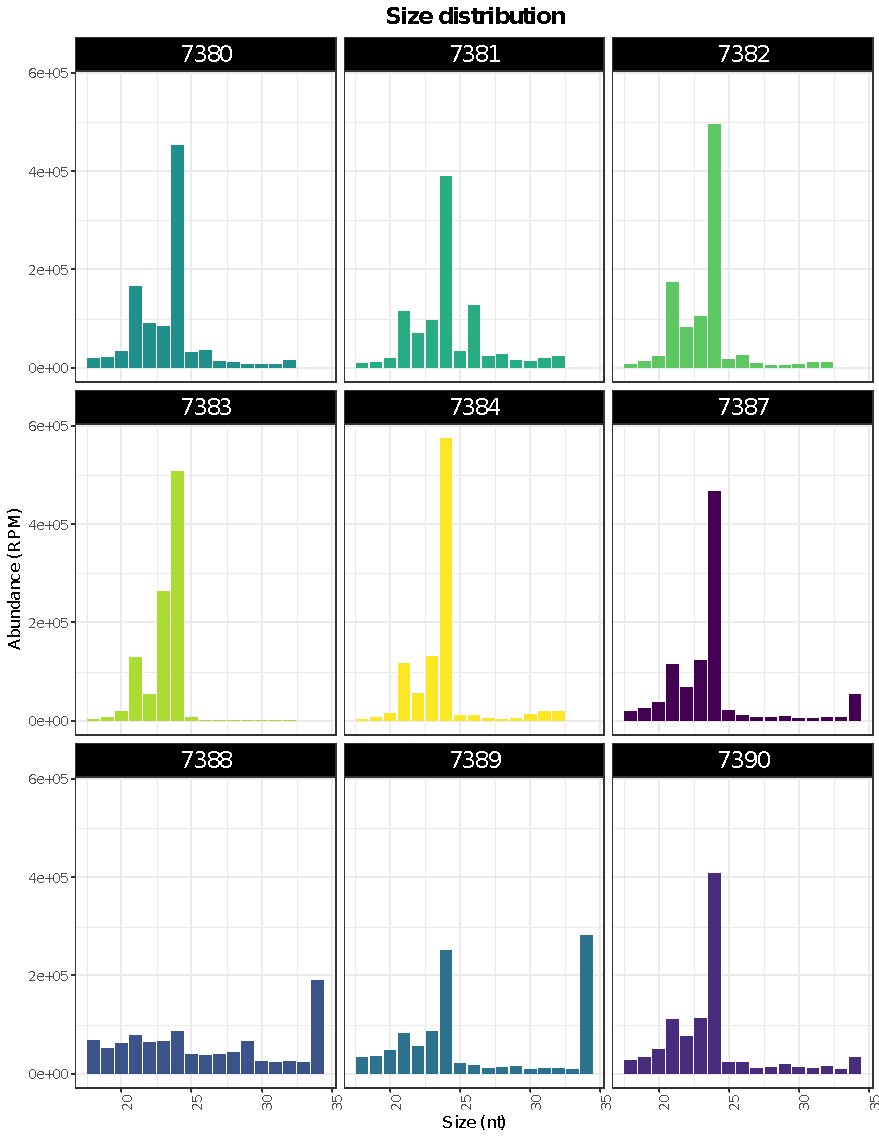
